# Compartmentalization of the SUMO/RNF4 pathway by SLX4 drives DNA repair

**DOI:** 10.1101/2022.09.20.508711

**Authors:** Emile Alghoul, Matteo Paloni, Arato Takedachi, Serge Urbach, Alessandro Barducci, Pierre-Henri Gaillard, Jihane Basbous, Angelos Constantinou

## Abstract

SLX4, disabled in Fanconi anemia group P, is a scaffolding protein that coordinates the action of structure-specific endonucleases and other proteins involved in replication-coupled repair of DNA interstrand crosslinks (ICLs). Here we show that SLX4 dimerization and SUMO-SIM interactions drive the assembly of SLX4 membraneless compartments in the nucleus called condensates. Super-resolution microscopy reveals that SLX4 forms chromatin-bound clusters of nanocondensates. We report that SLX4 compartmentalizes the SUMO-RNF4 signaling pathway. SENP6 and RNF4 regulate the assembly and disassembly of SLX4 condensates, respectively. SLX4 condensation *per se* triggers the selective modification of proteins by SUMO and ubiquitin. Specifically, SLX4 condensation induces ubiquitylation and chromatin extraction of topoisomerase 1 DNA-protein cross-links. SLX4 condensation also induces the nucleolytic degradation of newly replicated DNA. We propose that the compartmentalization of proteins by SLX4 through site-specific interactions ensures the spatiotemporal control of protein modifications and nucleolytic reactions during DNA repair.

## Introduction

In response to DNA damage, hundreds to thousands of copies of DNA damage response (DDR) proteins concentrate within nuclear foci ^1–4^. The assembly of DDR foci is governed by a network of site-specific interactions, as revealed by interdependencies in protein recruitment at DNA damage sites ^1, 3^. Such mesoscale structures that concentrate molecules in the absence of a surrounding membrane and no fixed stoichiometries are referred to as biomolecular condensates^5–11^, a term that makes no assumptions about the mechanism by which these structures are generated ^8, 12^. Several condensation mechanisms have been proposed, including liquid-liquid phase separation ^5, 8, 10, 13^, surface condensation ^14, 15^, polymer-polymer phase separation ^16, 17^, head-to-tail polymerization ^18–21^, and local protein enrichment at clustered binding sites^17, 22, 23^. Central to condensates biogenesis are multivalent and cooperative interactions involving folded protein and/or nucleic acid binding domains and intrinsically disordered regions ^5, 8, 24–29^.

Protein clustering is typically driven by a few key multivalent scaffolds that are highly connected to other molecules ^25, 26, 28^. A localization-induction model suggests that posttranslational modifications that increase attractive interactions trigger a transition towards the formation of stimuli-responsive condensates^30^. Of note, protein group modification by SUMO stabilizes protein interactions enhancing biochemical reactions, as exemplified in homologous recombination and ATR activation ^31, 32^. Likewise, SUMO-SIM interactions stabilize Promyelocytic leukemia protein (PML) nuclear bodies ^33–36^. Consistent with this, synthetic poly(SUMO) and poly(SIM) scaffolds self-assemble and form condensates *in vitro* ^37^. Client proteins can be recruited to condensates by binding to the same interacting domain involved in the condensation of the scaffolds^37^. Compartmentalization of the SUMOylation machinery within a synthetic condensate markedly enhances the rates of protein SUMOylation ^38^, suggesting that the organization of enzymes and substrates within condensates facilitates biochemical reactions. However, the function of natural cellular condensates in controlling protein SUMOylation rates and specificity remains to be demonstrated.

Here we report on the low abundance protein SLX4, a scaffold required for genome integrity. Biallelic inactivation of *SLX4* underlies complementation group P of Fanconi anemia ^39, 40^, an inherited disease associated with congenital malformations, pancytopenia and cancer susceptibility ^41^. SLX4 associates with several DNA repair factors including the structure-specific endonucleases (SSEs) XPF-ERCC1, MUS81-EME1, and SLX1 to repair DNA interstrand crosslinks, resolve Holliday junctions, ensure telomere maintenance, and maintain stability of common fragile sites ^42–53^. Recent evidence indicates that at a replication fork barrier caused by tightly DNA-bound proteins, SLX4-XPF promotes the recruitment of DNA damage response factors and DNA repair by homologous recombination ^54, 55^. SLX4 and XPF deficient cells are hypersensitive to DNA methyltransferases and topoisomerase 1 DNA-protein crosslinks (DPCs) ^54, 56, 57^, but how SLX4-XPF mechanistically promotes the processing of DPCs remains incompletely understood.

We show that SLX4 assembles nuclear condensates that are held together by a structurally defined oligomerization domain and site-specific SUMO-SIM interactions. Control of SLX4 condensation with high temporal precision revealed that SLX4 activates the SUMO/RNF4 pathway and ensures the selective modification of substrate proteins through the formation of membraneless compartments. SLX4 condensation significantly enhanced the extraction of topoisomerase 1 DNA-protein crosslinks (DPCs) from chromatin, unmasking the direct contribution of SLX4 to the removal of tightly DNA-bound proteins. Furthermore, SLX4 condensation induced the degradation of nascent DNA in the absence of exogenous sources of DNA damage. We propose that SLX4 promotes and controls by compartmentalization a cascade of enzymatic reactions determined by the composition of SLX4 condensates.

## Results

### SLX4 is a scaffolding component of nuclear foci

We took advantage of a cryptochrome 2 (Cry2) based optogenetic tool to mechanistically dissect the behavior of SLX4 in living cells and used mCherry for direct visualization of Cry2-SLX4, called optoSLX4 thereafter (Figure 1A). Cry2 forms tetramers upon exposure to 488 nm light ^58^. This optogenetic module can be used to modulate protein interactions in cells^59^. When fused to a multivalent protein scaffold underlying nuclear foci, Cry2 oligomerization triggers the scaffold’s condensation, providing unprecedented spatiotemporal control over the formation of DDR foci, in the absence of DNA damage ^6, 7^. We controlled the expression of the recombinant optoSLX4 protein with doxycycline in HEK293 Flp-In T-Rex cells (Figure 1A) and performed optogenetic activation using four-second exposures to 488 nm light, separated by 10-second intervals, for a total period of three minutes. This protocol induced the formation of multiple SLX4 foci per nucleus (Figure 1B, C). Optogenetic SLX4 foci were detectable as early as 30 seconds after blue light exposure and occasionally clustered (Figure 1D, Video S1, S2, S3). This indicates that SLX4 is a key determinant of nuclear foci formation.

**Figure 1.**
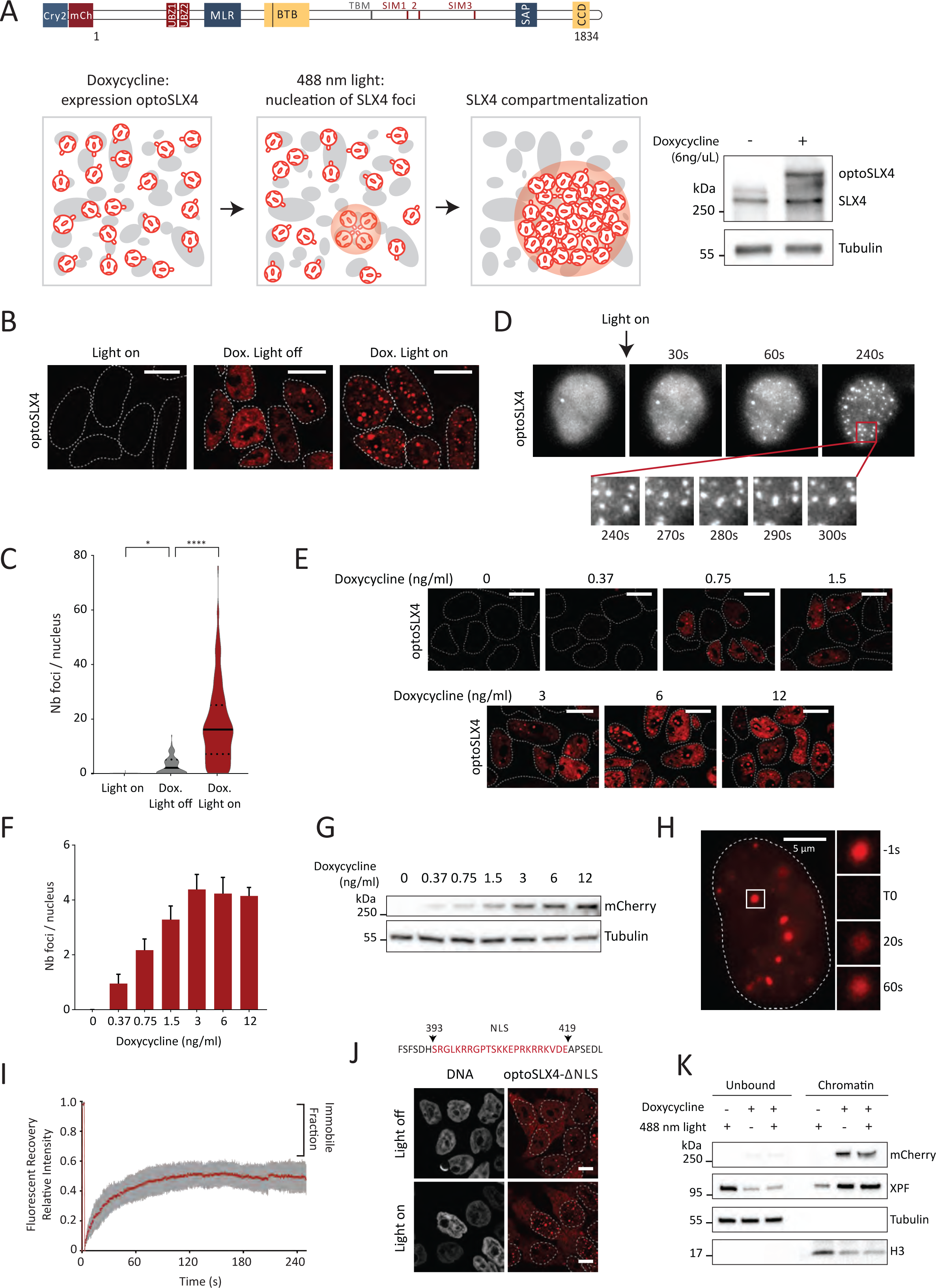
SLX4 drives the assembly of nuclear condensates. (A) Upper panel, schematic of the optoSLX4 construct: Cry2: cryptochrome 2, mCh: mCherry. Lower left, schematic representation of light induced SX4 condensation. Lower right, immunoblotting for SLX4. (B) Representative images of optoSLX4 condensates. Doxycycline (Dox.). Scale bar: 10 μm. (C) Violin plot quantification of optoSLX4 condensates. Median (plain line), quartile (dashed line); n=2, >100 cells per condition, *p < 0.05, ****p < 0.0001. (D) Representative time-lapse microscopy images of light induced SLX4 condensation and fusion events. Time: seconds (s). (E) Representative images of optoSLX4 expressed with the indicated concentration of doxycycline Scale bar: 10 μm. (F) Quantitative histogram analysis of spontaneous SLX4 condensates. Data plotted as medians with SEM; n= 3, >100 cells per condition. (G) Immunoblotting with the indicated antibodies shows SLX4 expression level in (E) and (F). (H) Representative images of a FRAP experiment. Scale bar 5 μm. (I) Quantifications of the FRAP data in (H) shown as mean ± SEM (n = 10). (J) Upper panel, NLS sequence deleted. Lower panel: representative images of light induced SLX4 condensates. Scale bar 10 μm. (K) Immunoblotting with the indicated antibodies of fractionated cell extracts prepared from cells treated with doxycycline and light when specified with (+).

Since protein condensation is a concentration-dependent process, we examined the relationship between optoSLX4 condensation and optoSLX4 expression levels. The number of spontaneous SLX4 condensates was directly proportional to the level of SLX4 expression induced by doxycycline, until a plateau was reached above 3ng/ml doxycycline (Figure 1E, F). An mCherry SLX4 construct lacking Cry2 confirmed that in the absence of blue-light, Cry2 does not play a role in the concentration-dependent assembly of spontaneous SLX4 condensates (Figure S1E-G).

DDR foci are reversible structures that disassemble upon completion of the repair process. To verify that optogenetic activation of SLX4 produces reversible condensates, as opposed to solid aggregates, we examined the timing of their dissolution. OptoSLX4 condensates dissolved progressively 20 to 30 minutes after optogenetic activation (Figure S1A, B). The lifetime of Cry2 interactions is approximately 1-2 minutes in the absence of light ^59^, so the lifetime of optogenetic SLX4 condensates mainly reflects the contribution of SLX4 interactions to the stability of these mesoscale structures. This behavior is similar to that of optogenetic TopBP1 condensates ^6^. To compare the dynamics of SLX4 condensates induced by optogenetic activation with the dynamics of SLX4 foci induced by a chemotherapeutic agent that causes DNA damage, we exposed optoSLX4-expressing cells to camptothecin (CPT), an inhibitor of topoisomerase 1 (TOP1). SLX4 is required for cellular tolerance to CPT ^56, 57^. TOP1 resolves DNA supercoils by introducing a transient nick into the DNA. During this process, TOP1 forms a transient covalent reaction intermediate with DNA called the TOP1 cleavage complex (TOP1cc) ^60^. Endogenous DNA lesions or specific inhibitors of TOP1 such as CPT trap TOP1cc, resulting in topoisomerase 1 DNA-protein crosslinking (TOP1-DPC) ^60^. CPT induced the accumulation of SLX4 foci during the course of a one-hour incubation (Figure S1C, D). Upon removal of CPT, SLX4 foci almost completely disappeared after 30 minutes (Figure S1C, D). The data indicate that SLX4 foci induced by CPT or optogenetic activation exhibit similar dissolution kinetics.

To gain further insights into the nuclear dynamics of optoSLX4 condensates, we examined fluorescence recovery after photobleaching. Two minutes after SLX4 foci photobeaching, fluorescence recovery was approximately 50%, suggesting that SLX4 foci contain a high mobility and a low mobility population of SLX4 molecules (Figure 1H, I), consistent with the dynamics of proteins within condensates formed on chromatin ^61–64^. To investigate the contribution of the subcellular environment to the formation of optogenetic SLX4 condensates, we deleted a predicted bipartite nuclear localization signal (NLS) in SLX4 (Figure 1J). We detected optoSLX4 - ΔNLS in both the cytoplasm and nucleoplasm. An additional putative NLS such as the NLS predicted in the 118 to 129 amino acid sequence may be responsible for the residual nuclear localization of SLX4. The cellular behavior of optoSLX4 - ΔNLS provided us with the opportunity to test the propensity of SLX4 to assemble condensates in these two subcellular environments. Remarkably, optogenetic SLX4-ΔNLS condensates formed exclusively in nuclei (Figure 1J). SLX4 signals remained diffuse in the cytoplasm (Figure 1J), suggesting that the assembly of SLX4 condensates is driven by modifications and interactions that are specific to the nucleoplasmic environment. To verify that optoSLX4 associates with chromatin, we fractionated cells into chromatin and soluble fractions (Figure 1K). We detected chromatin-bound optoSLX4 both in untreated cells and in cells exposed to blue light (Figure 1K). Induction of optoSLX4 expression with doxycycline decreased the soluble fraction and increased the chromatin bound fraction of the SLX4-associated endonuclease XPF-ERCC1 (Figure 1K), consistent with the scaffolding function of SLX4. The data thus far indicate that SLX4 supports the assembly of membraneless compartments on chromatin, that the process of SLX4 compartmentalization is reversible and concentration dependent, and that SLX4 molecules in these compartments exchange with the surrounding nucleoplasm. These observations recapitulate characteristic features of biomolecular condensates.

We next performed co-localization analyses to gain further evidence that optogenetic SLX4 condensates mimic endogenous SLX4 foci. Light-induced SLX4 condensates co-localized with endogenous TopBP1, BRCA1, MDC1, TRF2, PML and RPA32 (Figure 2A). Quantification of overlapping fluorescence signals revealed a wide range of signal overlap, ranging from 10% to 80% (Figure 2B). Thus, the composition of SLX4 condensates was subject to changes, most likely during the cell cycle and depending on specific cellular cues. We quantified SLX4 foci induced by optogenetic activation throughout the cell cycle using the fluorescent ubiquitylation-based cell cycle indicator PIP-FUCCI ^65^. Light induced SLX4 foci were detected not only in the S and G2/M phases of the cell cycle, but also in the G1 phase, albeit to a lesser extent (Figure 2C, D), consistent with a previous report ^55^. These observations suggest that optogenetic SLX4 condensates recapitulate the assembly of SLX4 foci.

**Figure 2.**
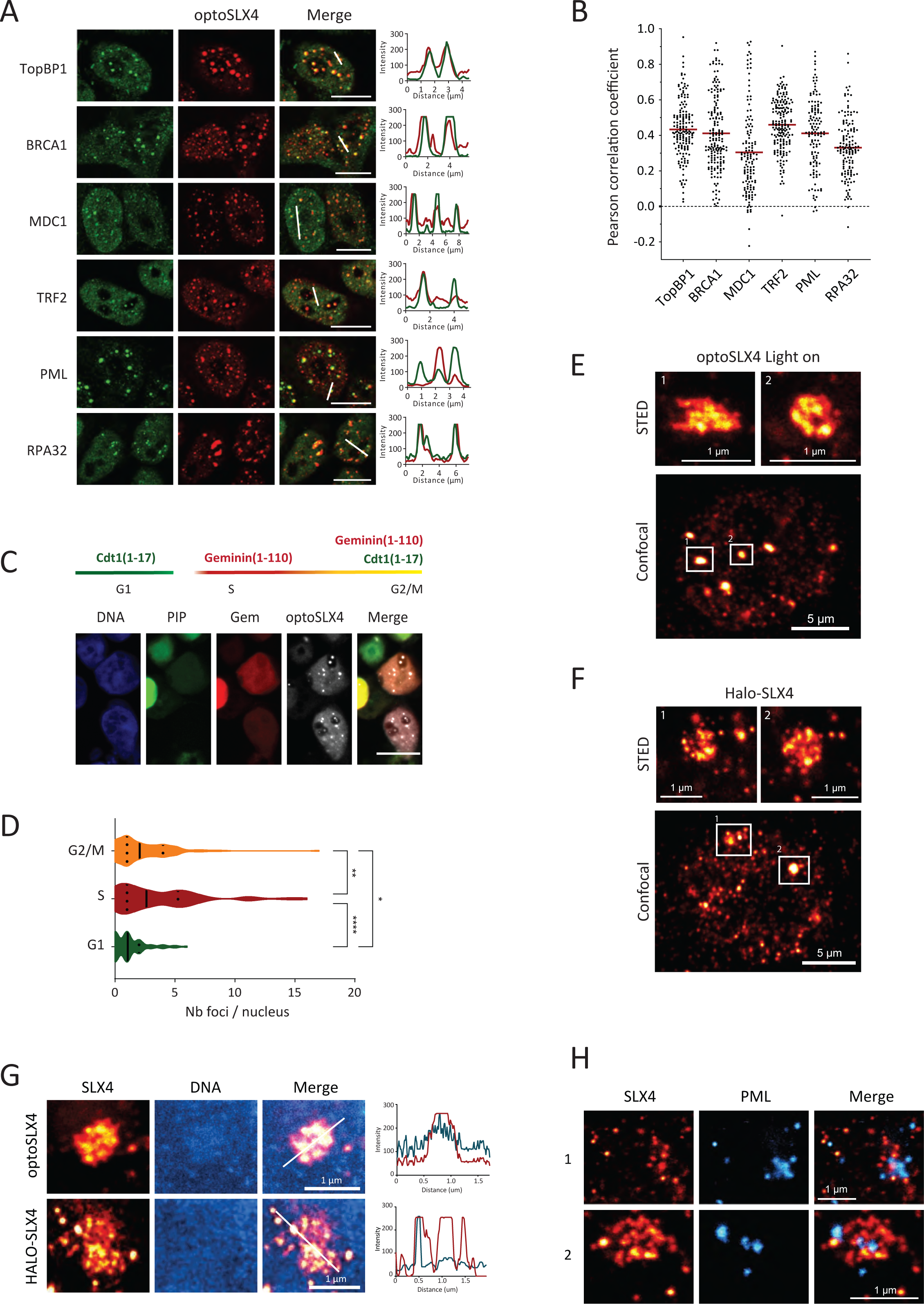
OptoSLX4 condensates recapitulate endogenous SLX4 foci. (A) Representative images of optoSLX4 condensates (red channel) and the indicated proteins revealed by immunofluorescence staining (green channel). Line scans of the red and green channels are shown. Scale bar 10 μm. (B) Quantification, using Pearson’s correlation coefficient, of the co-localization between the indicated proteins and optoSLX4 condensates. Median shown in red, >100 cells per condition. (C) Representative images of the distribution of optoSLX4 condensates throughout the cell cycles analyzed with the FUCCI cell cycle reporter. Scale bar 10 μm. (D) Violin plot quantification of optoSLX4 condensates in G2/M, S and G1, as indicated. Median: plain line, quartile: dashed lines. n=2, >100 cells, *p < 0.05, **p < 0.01, ****p < 0.0001. (E) Representative confocal and STED images of optoSLX4 condensates. (F) Representative confocal and STED images of endogenous Halo-SLX4 condensates. (G) Representative STED images of optoSLX4, endogenous Halo-SLX4 (red channel) and DNA (blue channel). Line scans of the red and blue channels are shown. (H) Representative STED images of SLX4 (red channel) and PML (blue channel).

Last, we used super-resolution imaging by stimulation-emission-depletion (STED) microscopy to gain insights into the intrinsic organization of SLX4 condensates. We observed that SLX4 forms globular clusters of nanocondensates of approximately 100 nm in size (Figure 2E, F), similar to clusters of γH2AX, 53BP1 and TopBP1 nanofoci ^6, 66–68^. DNA signals were enriched within SLX4 condensates, suggesting a compact structure (Figure 2G). To directly visualize endogenous SLX4 with fluorescent ligands, we tagged *SLX4* alleles with HaloTag. In comparison with Halo-SLX4-depleted cells, cells expressing Halo-tagged SLX4 were resistant to mitomycin C, confirming that the HaloTag preserves the function of SLX4 in DNA interstrand crosslink repair (Figure S2A-C). Remarkably, cells expressing physiological levels of SLX4 detected with HaloTag fluorescent ligands exhibited spontaneous SLX4 foci throughout the cell cycle (Figure S2D). Endogenous SLX4 nanocondensates formed globular clusters that were indistinguishable from recombinant optoSLX4 condensates (Figure 2F). Taken together, these observations suggest that optogenetic SLX4 condensates assembled on demand with blue light closely mimic endogenous SLX4 foci.

Parenthetically, we also analyzed the localization of PML relative to SLX4. While conventional fluorescence microscopy shows that PML and SLX4 partially colocalize (Figure 2A), super-resolution microscopy revealed that PML and SLX4 signals exhibit distinct spatial distribution (Figure 2H). PML was often proximal to SLX4 yet did not overlap with SLX4 nanocondensates (Figure 2H).

### Site-specific interactions drive the biogenesis of SLX4 condensates

Since our ultimate goal was to understand the functions of SLX4 condensation, it was first necessary to identify key determinants of SLX4 condensation amenable to structure-function analyses. SLX4 is largely unstructured with only a few folded domains ^69^. We generated a series of truncated SLX4 proteins fused to mcherry-Cry2 to gain insight into the domains involved in condensate assembly (Figure S2E-G). Optogenetic activation of the Bric-a-brac Tramtrack and Broad complex (BTB) oligomerization domain alone did not yield condensates (Figure S2E-G), but deletion of the BTB domain reduced the ability of SLX4 to assemble condensates (ΔBTB, Figure S2E-G), even though the protein was fused to Cry2, which forms tetramers upon light activation. Thus, the BTB domain contributes to SLX4 condensation. The intrinsically disordered amino terminal region of SLX4 (IDR1) did not form detectable condensates (Figure S2F), however, IDR1 was less stable than WT SLX4 (Figure S2G). IDR1 was more stable in combination with the BTB domain (Figure S2G), but the IDR1-BTB truncated protein also failed to form condensates (Figure S2F). In contrast, the intrinsically disordered carboxyl terminal portion of SLX4 (IDR2) showed partial ability to assemble SLX4 condensates (Figure S2F), and the fusion of IDR2 to the BTB further stimulated SLX4 condensation (Figure S2F). Thus, the BTB-IDR2 portion of SLX4 enables the formation of SLX4 condensates through optogenetic activation. IDR2 contains multiple SUMOylation sites as well as three SUMO-interacting motifs (SIMs), which can in principle promote the formation of biomolecular condensates ^37, 53, 56, 70, 71^.

To evaluate how the BTB and SIMs domains in SLX4 might contribute to the assembly of nuclear condensates, we first relied on molecular simulations. We modeled SLX4 monomer and BTB-mediated dimers as flexible chains of beads representing individual domains and connected by harmonic springs (Figure 3A), following a recent approach to simulate the condensation of associative biopolymers ^72^. In this framework, the specific protein - protein interaction between SUMO and SIM domains is modeled by an attractive potential while a strong repulsion between domains of the same type ensures a correct one-to-one binding stoichiometry (Figure S3A). Further details are described in the Methods section. Using this coarse-grained approach, we performed extensive simulations to explore condensate formation of SLX4 monomers or dimers with different degrees of SUMOylation (Figure 3B, C). While this minimal model necessarily underestimates the number and diversity of intermolecular interactions that occur in cells, this simulation shed some light on the molecular determinants of SLX4 assembly. Overall, we observed that BTB-mediated dimerization strongly enhanced SLX4 condensation independent of the total protein concentration due to the higher interaction valency of SLX4 dimers (Figure 3B, C, S3B). In this model, the degree of SUMOylation of SLX4 chains determines SLX4 assembly in a non-monotonic manner. While the presence of multiple SUMOylated sites significantly enhanced the formation of SLX4 condensates, this trend was partially reversed when most of the potential SLX4 SUMO sites were SUMOylated (Figure 3B, C, S3B). This behavior results from the effective saturation of the three available SIMs with SUMO groups from the same molecule. Thus, simulations suggest that the balance between intra- and intermolecular binding makes the condensation propensity dependent on the specific patterning of SUMO groups on SLX4 sites (Figure S3D, E). However, the model also suggests that efficient assembly of SLX4 condensates at nanomolar nuclear concentrations must rely on further intermolecular interactions, such as chromatin-SLX4 binding, which could increase the local concentration of SLX4.

**Figures 3.**
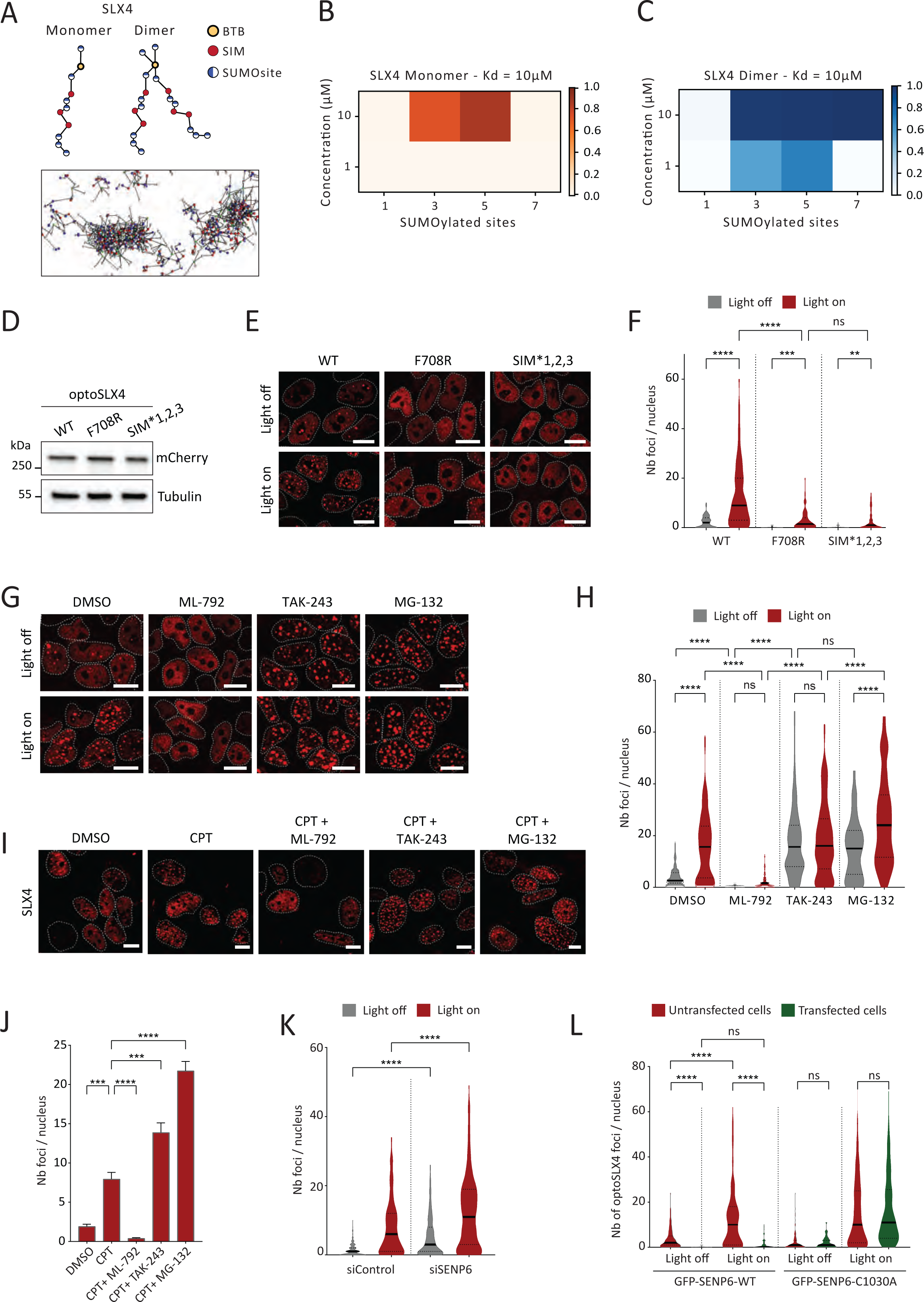
SLX4 oligomerization and SUMO-SIM interactions promote SLX4 condensation. (A) Upper panel: Schematic representation of the molecular models of SLX4 monomer (left) and BTB-mediated dimer (right). Yellow particles are BTB domains, red particles are SIM domains, blue and white particles are SUMOylation sites. (lower panel) Representative snapshot of condensate formation from a MD simulation of BTB-mediated dimers of SLX4 with three SUMOylated sites per monomer. Molecular models of SLX4 monomer (left) and BTB-mediated dimer (right). Lower panel: Representative snapshot of condensate formation (Dimers, three SUMOylated sites per monomer). (B) Fraction of chains in the condensed phase as a function of the total concentration and the number of SUMOylated sites per monomer in simulations of mixtures of SLX4 monomers, where the Kd for SUMO-SIM interaction is set to 10 μM. (C) Fraction of chains in the condensed phase as a function of the total concentration and the number of SUMOylated sites per monomer in simulations of mixtures of BTB-mediated SLX4 dimers, where the Kd for SUMO-SIM interaction is set to 10 μM. (D) Expression of optoSLX4 wt, and F708R and SIM *1,2,3 mutants revealed by immunoblotting. (E) Representative images before and after exposure to 488 nm light. Scale bar 10 μm. (F) Violin plot quantification of (E). Median (plain line) and quartile (dashed line) indicated. n= 3, >100 cells per condition, ns: non-significant, **p < 0.01, ***p < 0.001, ****p < 0.0001. (G) Representative images of optoSLX4 cells treated with ML-792 (2 μM), TAK-243 (2 μM) and MG-132 (10 μM) for 4 hr. Scale bar 10 μm. (H) Violin plot quantification of (G). Median (plain line) and quartile (dashed line) indicated. n= 2, >100 cells per condition, ns: non-significant, ****p < 0.0001. (I) Representative images of FA-P cells corrected with wtSLX4 treated with ML-792 (2 μM), TAK-243 (2 μM) and MG-132 (10 μM) for 4 hr plus CPT (1 μM) during the last hour. (J) Quantitative histogram analysis of SLX4 foci per nucleus (E). Data plotted as median with SEM; n= 2, >200 cells per condition, ***p < 0.001, ****p < 0.0001. (K) Violin plot quantification of optoSLX4 foci in cells treated with the indicated siRNAs before and after light activation. Median (plain line), quartile (dashed line). n= 2, >200 cells per condition, ****p < 0.0001. (L) Violin plot quantification of optoSLX4 foci in cells transfected with GFP-SENP6-WT or GFP-SENP6-C1030A, as indicated, before and after light activation. Median (plain line), quartile (dashed line). n= 2, >100 cells per condition, ns: non-significant, ****p < 0.0001.

Consistent with simulations, a phenyl to arginine substitution (F708R) in the BTB domain, which disrupts a key contact required for SLX4 oligomerization ^73^, impaired the optogenetic activation of SLX4 condensates (Figure 3D-F). In the absence of Cry2, the mCherry-SLX4-F708R mutant protein did not form concentration-dependent SLX4 foci (Figure S4A-C), indicating that the residual foci formed by the mCherry-Cry2-SLX4-F708R mutant protein were dependent on blue-light induced oligomerization of Cry2. This confirms that the SLX4 dimerization is required for SLX4 condensation. Substitution of all the aliphatic acids of SIM 1, 2 and 3 with alanine (SIM*1,2,3) also undermined SLX4 condensation (Figure 3D-F). Next, we induced osmotic stress with NaCl, sucrose or sorbitol to increase protein concentrations in the nucleoplasm ^74^. This treatment was sufficient to induce the formation of SLX4 foci, but the SLX4-F708R and SLX4-SIM*1,2,3 mutant proteins did not form foci under these experimental conditions (Figure S4D, E). Preincubation of cells with an inhibitor of the SUMO activating enzyme (SAE), ML-792, impaired the formation of optogenetic SLX4 foci (Figure 3G, H). In contrast, inhibition of the ubiquitin activating enzyme with TAK-243 or the proteasome with MG-132 stabilized SLX4 structures (Figure 3G, H), suggesting that a SUMO-ubiquitin circuit controls the self-assembly of SLX4 at the mesoscale, as discussed below.

Next, we used a different cellular system to confirm the role of SUMOylation and ubiquitylation in the regulation of SLX4 condensates. We exposed Fanconi anemia patient derived *SLX4* null cells (FA-P) complemented with WT *SLX4* cDNA to CPT for one hour and detected SLX4 foci by immunofluorescence staining. Preincubation of cells with ML-792 blocked the formation of SLX4 foci induced by CPT, whereas TAK-243 and MG-132 significantly increased the number of SLX4 foci (Figure 3I, J). Finally, the soluble fraction of the F708R SLX4 mutant protein was higher, and its chromatin bound fraction was decreased compared to wild-type optoSLX4 (Figure S4F). Mutations of the SIM motifs also increased the solubility of SLX4, as did the inhibition of SAE with ML-792 (Figure S4F). Taken together, these data suggest that the BTB oligomerization domain in combination with SUMO-SIM interactions promote the assembly of SLX4 condensates on chromatin.

The SUMO isopeptidase SENP6 directly regulates the size of PML nuclear bodies by acting upon the SUMO modified PML substrate protein ^33^. Similarly, the activity of SUMO proteases may antagonize the condensation of SLX4. Since SLX4 is a major target of SENP6 ^75, 76^, we depleted SENP6 using RNA interference to investigate its role in the regulation of SLX4 condensation (Figure S4G). Suppression of SENP6 increased the number of both spontaneous and light induced SLX4 foci per nucleus (Figures 3K and S4H). By contrast, spontaneous and light induced SLX4 foci were reduced in cells transfected with recombinant GFP tagged WT SENP6, whereas transfection of a catalytically dead mutant GFP-SENP6-C1030A had no effect SLX4 condensation (Figures 3L, S4I). GFP-SENP6-C1030A co-localized with optogenetic SLX4 foci, indicating that SENP6 was recruited to these compartments (Figure S4I). Thus, SUMO-SIM interactions promote the assembly of SLX4 condensates under the control of SENP6.

### SLX4 compartmentalizes the SUMOylation/Ubiquitylation system

Biomolecular condensates act as membraneless compartments that concentrate proteins selectively. To gain insights into the composition of SLX4 condensates, we used a biotin proximity labeling approach coupled to mass spectrometry ^77^. We reproducibly identified SLX4-associated proteins, including known SLX4 interactors such as XPF, MUS81, SLX4IP and TopBP1. We also identified SUMO2/3 and the E3 SUMO ligases RanBP2, PIAS1 and PIAS4 (Figure 4A), as well as ubiquitin and the E3 ubiquitin ligases TRIM25 and TRIM33, consistent with a recent proteomic study ^78^ . Since SLX4 is an essential component of a multifactorial SUMO E3 ligase that remains to be defined ^53^, we hypothesized that SLX4 may compartmentalize SUMO and ubiquitin modifying enzymes.

**Figure 4.**
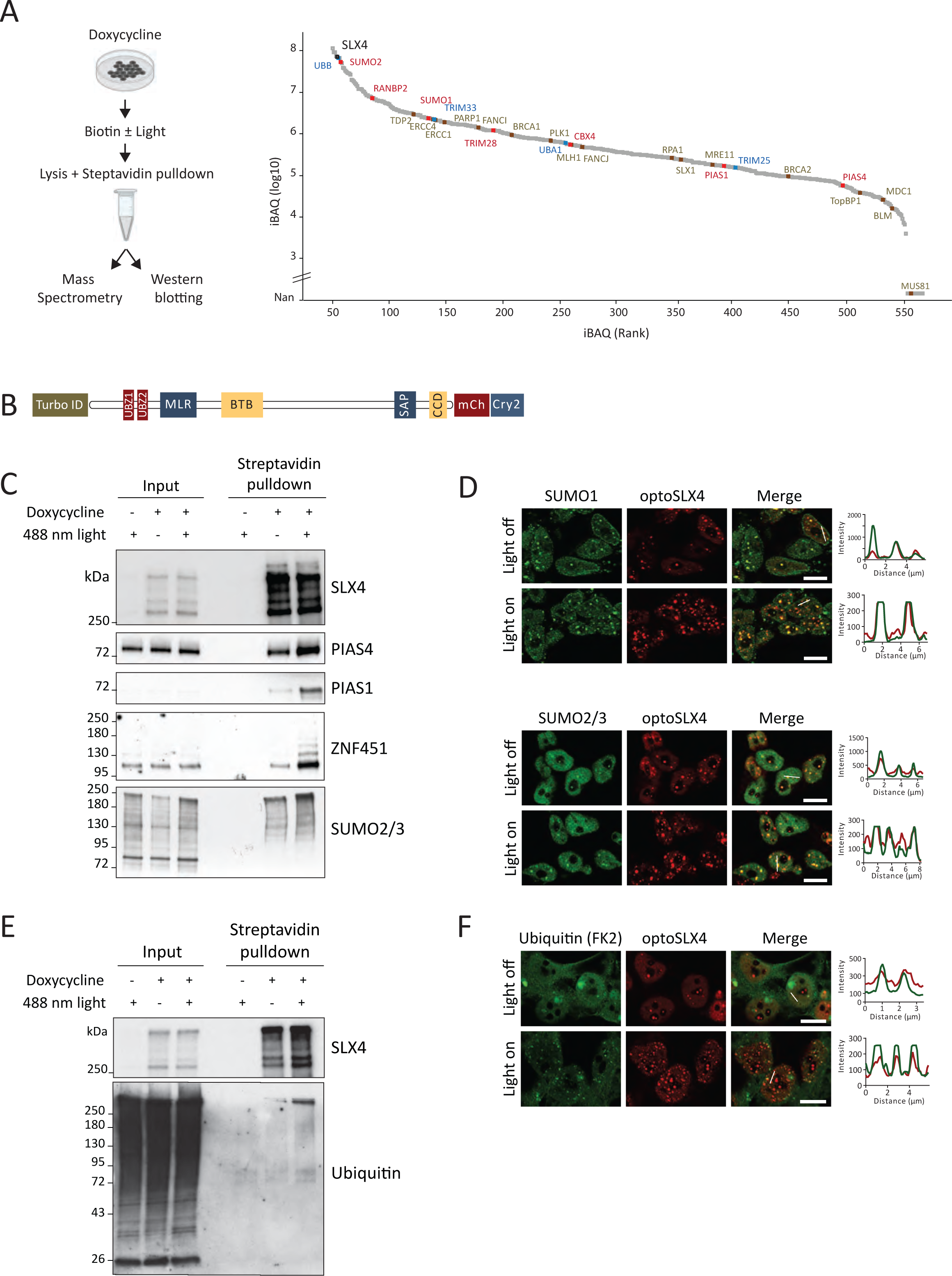
SLX4 condensates compartmentalize the SUMO/ubiquitin machinery. (A) Left, schematic of the proximity labeling experiments. Right, proteins identified by mass spectrometry and ranked according to their iBAQ values. The values correspond to the sum of iBAQs from seven independent experiments. Red: components of the SUMOylation machinery, blue: components of the ubiquitylation machinery, beige: examples of known SLX4 interactors. (B) Schematic of the SLX4 construct used for optogenetic (mCh/Cry2 module) coupled biotin proximity labeling (TurboID) experiments. (C) Immunoblotting for the indicated proteins isolated with streptavidin beads from optoSLX4 cells treated with doxycycline and exposed to 488 nm light, when specified (+). Biotin was added to the media in all conditions. (D) Representative immunofluorescence images of optoSLX4 (red channel), endogenous SUMO1 and SUMO2/3 (green channel). Line scans on the right show co-localization. Scale bar 10 μm. (E) Immunoblotting for SLX4 and ubiquitinated proteins isolated with streptavidin beads from cells treated with doxycycline and exposed to 488 nm light, when indicated (+). Biotin was added to the media in all conditions. (F) Representative immunofluorescence images of optoSLX4 (red channel) and mono/poly-ubiquitin (green channel). Line scans on the right show co-localization. Scale bar 10 μm.

To test whether SLX4 condensates concentrate E3 SUMO ligases, we combined the optogenetic and the biotin proximity labeling approaches. We fused SLX4 to TurboID at its amino-terminus and to mcherry-Cry2 at its carboxyl terminus (Figure 4B). We exposed cells to blue light for 15 minutes with 4s light-30s rest cycles in the presence of biotin in the cell culture medium, as described ^79^. Cells were then lysed and biotin-labelled proteins were isolated using streptavidin-coated beads. Compartmentalization of SLX4 by light significantly increased the amount of the E3 SUMO ligases PIAS1, PIAS4 and ZNF451 and of SUMO2/3 labeled in the vicinity of SLX4 (Figure 4C), indicating that SUMO2/3 and the E3 SUMO ligases accumulate locally within SLX4 condensates. Immunofluorescence staining confirmed that endogenous and GFP-tagged SUMO1 and SUMO2/3 as well as GFP-UBC9 and GFP-PIAS4 colocalized with SLX4 condensates (Figure 4D, S5A).

To begin to investigate the functions that arise specifically from the condensation of SLX4, we evaluated the impact of SLX4 condensation on the conjugation of substrate proteins to SUMO. To achieve this, we transfected optoSLX4 expressing cells with His-SUMO2/3, induced SLX4 condensation with blue-light in the absence of DNA damage, lysed the cells under denaturing conditions and isolated proteins conjugated to SUMO by metal affinity purification (Figure 5A). Previous studies have shown that SLX4, XPF, MDC1 and BRCA1 are SUMOylated and that these modifications are important for the DNA damage response ^53, 80, 81^. We detected SUMOylated optoSLX4 and XPF one minute after optogenetic induction of SLX4 condensates, and the signals increased thereafter (Figure 5B). The SUMOylation of SLX4 and XPF depended on the integrity of the BTB domain and the SUMO interaction motifs of SLX4 (Figure 5C), which are necessary for SLX4 condensation. Likewise, SLX4 condensation promoted the SUMOylation of MDC1 and BRCA1 (Figure 5D). Furthermore, using a candidate approach, we identified endogenous EME1 as a SUMOylation substrate under the control of SLX4 condensation (Figure 5D). SLX4-driven SUMOylation of EME1 was further supported by ex vivo/in vitro SUMOylation assays^53^, where endogenous EME1 co-immunoprecipitated with recombinant SLX4 underwent extensive SUMOylation *in vitro* (Figure S5B).

**Figure 5.**
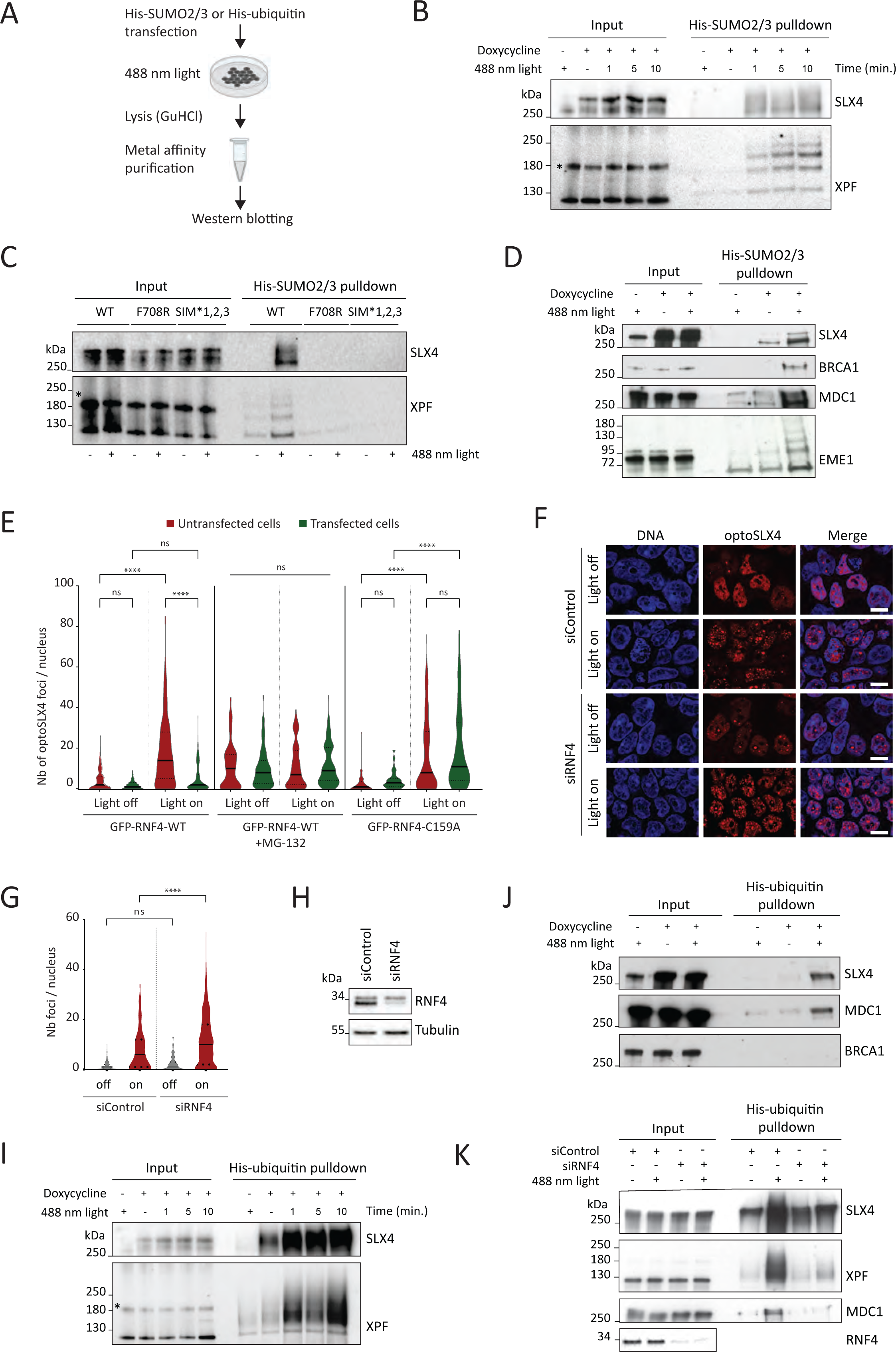
SLX4 condensation enhances the SUMOylation and ubiquitylation of substrate proteins. (A) Schematic of the experimental strategy. (B) Immunoblotting for SLX4 and XPF isolated via metal affinity pulldowns of proteins conjugated to His-SUMO2/3. Exposure times to 488 nm light is indicated in minutes (min.) (C) Isolation of SUMOylated proteins, as described in (A), from cells expressing wt, F708R and SIM*1,2,3 optoSLX4 proteins. Light (488nm) activation (+): 15 minutes. (D) Immunoblotting for the indicated proteins isolated via metal affinity pulldowns of His-SUMO2/3 conjugates. (E) Violin plot quantification of optoSLX4 foci in cells transfected with GFP-RNF4-WT or GFP-RNF4-C159A, as indicated, before and after light activation. When specified, cells were treated with MG132 (10μM) for 4 hr. Median (plain line), quartile (dashed line). n= 3, >100 cells per condition, ns: non-significant, ****p < 0.0001. (F) Representative images of optoSLX4 cells transfected with the indicated siRNAs. Scale bar 10 μm. (G) Violin plot quantification of (F). Median (plain line), quartile (dashed line). n= 2, >100 cells per condition, ns: non-significant, ****p < 0.0001. (H) Immunoblotting for the indicated proteins from cells treated with siControl and siRNF4. (I) Immunoblotting for XPF and SLX4 isolated via metal affinity pulldowns of His-ubiquitin conjugates. (J) Immunoblotting for the indicated proteins isolated via metal affinity pulldowns of His-ubiquitin conjugates. (K) Immunoblotting for the indicated proteins isolated via metal affinity pulldowns of His-ubiquitin conjugates from cells treated with siControl or siRNF4.

Protein SUMOylation plays important roles in the DDR, including the regulation of protein localization and the stabilization of physical interactions ^31, 82–84^. In addition, SUMO-modified proteins can be targeted for ubiquitylation and subsequent proteasomal degradation to accelerate protein turnover at DNA damage sites and enable DNA repair ^83, 85, 86^. To test whether SLX4 compartmentalizes the SUMO targeted ubiquitin ligase (STUbL) RNF4, we transfected cells with GFP tagged WT RNF4 or the ubiquitin ligase dead mutant GFP-RNF4-C159A. Accumulation of GFP-RNF4-WT within spontaneous and optogenetic SLX4 foci was only observed in the presence of the proteasome inhibitor MG-132 (Figure 5E, S5C). In contrast, the ubiquitin ligase dead mutant GFP-RNF4-C159A accumulated in SLX4 condensates even in the absence of proteasome inhibition (Figure 5E, S5C). Remarkably, transfection of GFP-RNF4-WT in the absence of proteasome inhibition instead significantly reduced the number of optogenetic SLX4 foci (Figure 5E, S5C). This observation suggests that RNF4-mediated protein ubiquitylation promotes the dissolution of SLX4 condensates. Consistent with this, the depletion of RNF4 by RNA interference increased the yield of light induced SLX4 foci (Figure 5F-H). Furthermore, SLX4 condensation increased the amount of ubiquitylated proteins detected in the vicinity of SLX4 (Figure 4E), and ubiquitin co-localized with SLX4 foci (Figure 4F). More specifically, SLX4 condensation readily induced robust ubiquitylation of SLX4 and XPF, as revealed by the immunodetection of proteins purified by metal affinity from cells transfected with His-ubiquitin (Figure 5I). Thus, condensation dependent SLX4 ubiquitylation represents a negative feedback regulatory mechanism leading to the dissolution of SLX4 condensates.

SLX4 compartmentalization also induced ubiquitylation of MDC1 (Figure 5J). In contrast, BRCA1 was not ubiquitylated, consistent with the finding that BRCA1 SUMOylation stimulates its E3 ubiquitin ligase activity ^81^, rather than its degradation by the proteasome. Finally, depletion of RNF4 by RNA interference inhibited the ubiquitylation of SLX4, XPF and MDC1 (Figure 5K). Taken together, these data indicate that SLX4 compartmentalizes and enhances the activity of the E3 SUMO ligases and STUbL RNF4 to selectively modify substrate proteins.

### SLX4 condensates enhance the degradation of topoisomerase 1 DNA - protein crosslinks

SUMO-targeted protein ubiquitylation and proteasomal degradation is emerging as a pathway for the degradation of DNA-protein crosslinks (DPCs), as illustrated by the degradation of TOP1-DPCs ^87^, of crosslinked DNA (cytosine-5)-methyltransferase 1 (DNMT1) ^88^, and of trapped PARP1 ^89^. TOP1-DPCs are modified by the PIAS4/RNF4 system and then degraded by proteolysis ^87^. This pathway is conserved in yeast where the human ortholog of RNF4 is the heterodimer Slx5-Slx8, ^87, 90^. Given the role of SLX4 in the compartmentalization of the SUMO/ubiquitylation system described above, the genetic relationship between *Slx4* and the *Slx5*-*Slx8* genes ^91^, and the hypersensitivity of SLX4-deficient cells to CPT^45, 46, 57^, we investigated the contribution of SLX4 condensation to the extraction of TOP1-DPCs ^87, 92^. SLX4 condensation significantly enhanced TOP1 conjugation to SUMO and ubiquitin (Figure 6A, upper and middle panels), and depletion of the STUbL RNF4 by RNA interference impaired TOP1 ubiquitylation within SLX4 compartments (Figure 6A, lower panel). We then treated cells with CPT, lysed the cells under strong denaturing conditions to disrupt non-covalent interactions, purified DNA along with crosslinked proteins, and examined TOP1-DPCs by immunoblotting. We isolated TOP1-DPCs from cells treated with CPT for 15 minutes (Figure 6B, C). Optogenetic activation of SLX4 condensates with 488 nm light pulses at the 5-minute and 10-minute time points during CPT treatment reduced the amount of TOP1-DPCs isolated from these cells (Figure 6B, C). Furthermore, immunodetection of TOP1-DPCs by fluorescence microscopy revealed high levels of TOP1-DPCs in Fanconi anemia patient derived *SLX4* null cells exposed to CPT (Figure 6D, E). Complementation of these cells with WT *SLX4* cDNA significantly reduced the level of TOP1-DPCs signals (Figure 6D, E). Conversely, the targeted degradation of the endogenous SLX4 HaloTag fusion protein mediated by the small molecule HaloPROTAC3 increased the levels of TOP1-DPCs in cells exposed to CPT (Figure 6F, G), as well as the cell intrinsic levels of TOP1-DPCs (Figure 6H, S5D). In contrast, doxycycline induced expression of recombinant SLX4 in HEK293 cells treated with CPT reduced TOP1-DPCs signals (Figure S5E). Taken together, these data indicate that compartmentalization of the SUMO/RNF4 pathway by SLX4 promotes the extraction of TOP1-DPCs from chromatin.

**Figure 6.**
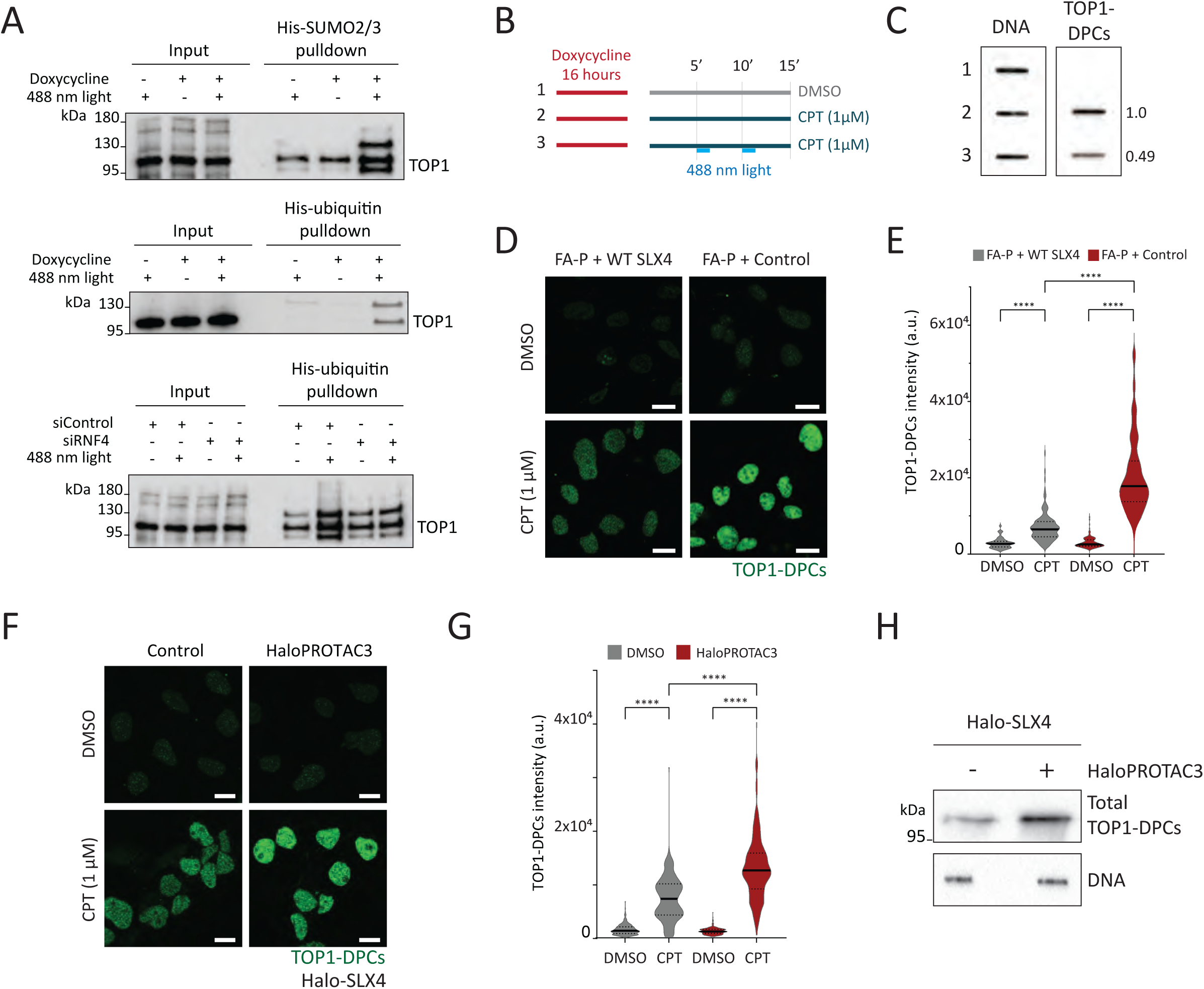
SLX4 condensation promotes removal of TOP1-DPCs. (A) Immunoblotting for TOP1 isolated after SLX4 condensation in pulldowns of His-SUMO2/3 conjugates (Upper panel), His-ubiquitin conjugates (middle panel), and His-ubiquitin conjugates from cells treated with siControl and siRNF4, as indicated (lower panel). (B) Schematic of the experiment. When indicated, light activation was performed for 1 minute during the 5th and 10th minute of the CPT treatment (15 minutes, 1 μM). (C) Slot blot of TOP1-DPCs and DNA, purified under chaotropic conditions following treatments described in (B). The DNA was used as a loading control. The relative amount of TOP1-DPCs is calculated by ImageJ. (D) Representative immunofluorescence images of TOP1-DPCs from FA-P cells and their corrected counterparts exposed to 1 μM CPT, as indicated. Scale bar: 10 μm. (E) Violin plot quantification of TOP1-DPCs intensity in (D). Median (plain line), quartile (dashed line). n= 3, >200 cells per condition, ****p < 0.0001. (F) Representative immunofluorescence images of TOP1-DPCs from Halo-SLX4 cells treated with HaloPROTAC3 and exposed to 1 μM CPT, as indicated. Scale bar: 10 μm. (G) Violin plot quantification of TOP1-DPCs intensity in (F). Median (plain line), quartile (dashed line). n= 3, >200 cells per condition, ****p < 0.0001. (H) Immunoblotting for TOP1-DPCs and DNA purified under chaotropic conditions. When indicated (+), cells were treated with HaloPROTAC3. DNA was used as a loading control.

### SLX4 condensation induces the degradation of newly replicated DNA

The data so far suggest SLX4 condensation enhances the SUMOylation and SUMO-dependent ubiquitylation of substrate proteins. Given that this system likely amplifies endogenous processes that would normally occur under highly regulated conditions, we investigated the consequences of SLX4 condensation on replication fork collapse, a process that normally occurs as a last resort mechanism when obstacles to replication fork progression persist ^53, 55, 93, 94^. In mammalian cells, the collapse of replication forks is driven by the SUMOylation and SUMO-targeted ubiquitylation of replisome components in a manner that depends on the combined action of RNF4, the AURKA-PLK1 pathway and SLX4 ^95^. We used a DNA fiber labeling approach to directly assess the effect of SLX4 condensation on DNA replication. We labeled cells with two consecutive pulses of iododeoxyuridine (IdU) and chlorodeoxyuridine (CldU) for 30 min each, and then exposed the cells to blue light for 3 min every 30 min for 2 h (Figure 7A). Since optogenetic SLX4 compartments spontaneously dissolve 30 minutes after induction (Figure S1A, B), this experimental procedure ensures the presence of SLX4 condensates for 2 hours. Replication forks proceed normally at a constant speed, hence the ratio of CldU/IdU replication track lengths is close to one (Figure 7B). However, after optogenetic activation of SLX4 condensates, the length of CldU replication tracks was shorter than that of IdU replication tracks (Figure 7B). In contrast, the ratio of CldU/IdU replication tracks remained close to 1 in cells expressing the oligomerization or SUMO interaction defective mutants F708R SLX4 and SIM*1,2,3, respectively (Figure 7B). Consistent with this, the ratio of CldU/IdU replication tracks was unaffected by 488 nm light in cells treated with the SAE inhibitor ML-792 (Figure 7C). These data suggest that in the absence of DNA replication inhibitors, SLX4 condensation either promotes the degradation of CldU-labeled DNA or induces the stalling of DNA replication forks. However, depletion of RNF4 by RNA interference blocked the shortening of CldU-labeled replication tracks (Figure 7D).

**Figure 7.**
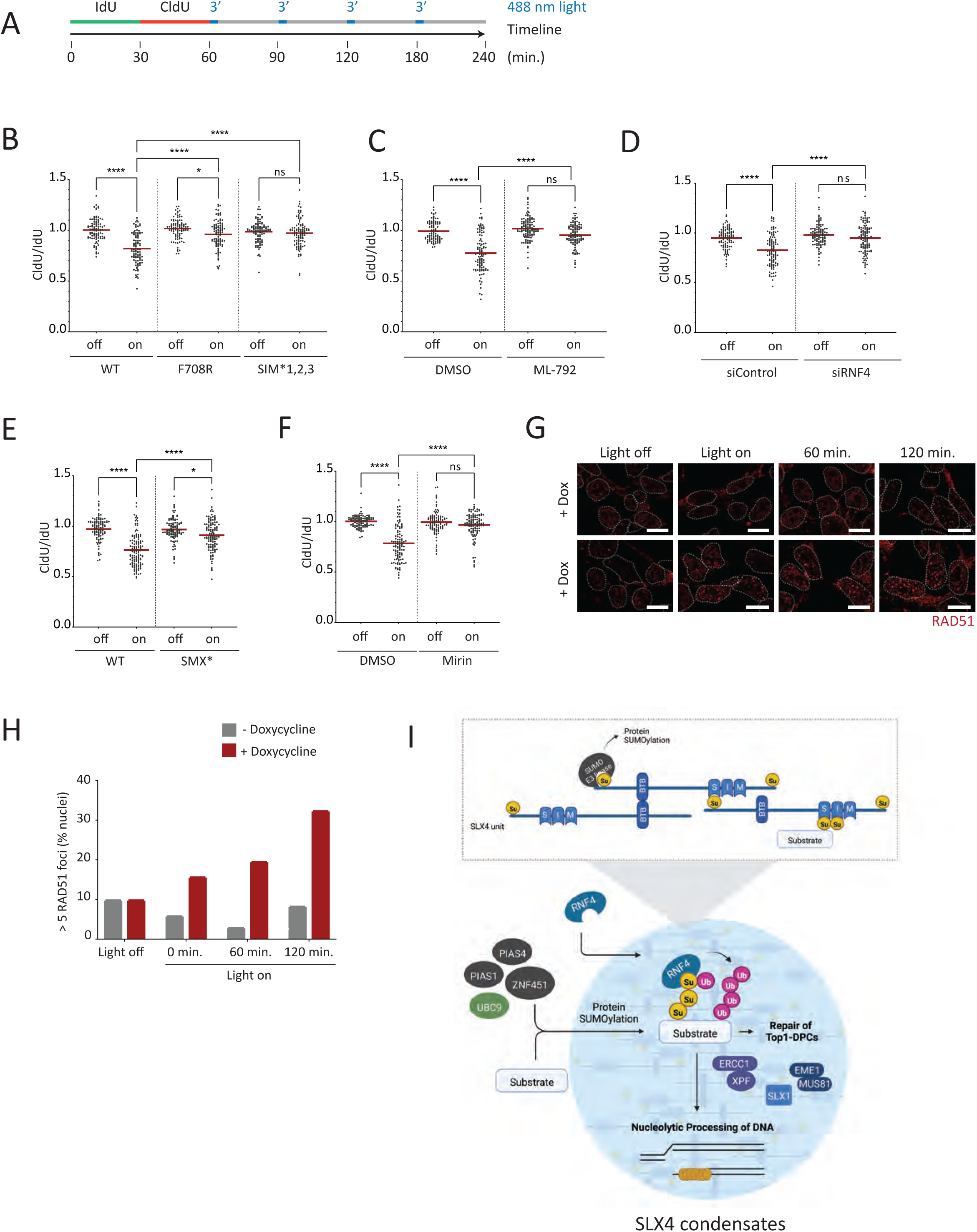
SLX4 condensation triggers nascent DNA degradation. (A) Schematic of the DNA fiber labelling experiment. (B) Dot plots represent CldU/IdU incorporation ratio of cells expressing WT, F708R and SIM*1,2,3 optoSLX4 in absence (off) or presence (on) of the light activation scheme described in (A). Mean values (red lines). n=3, > 100 fibers per condition, ns: non-significant, *p < 0.05, ****p < 0.0001. (C) CldU/IdU incorporation ratio from optoSLX4-WT cells pre-treated with the SAE inhibitor ML-792 (2μM) for 2 hr. Mean values (red lines). n=2, > 100 fibers per condition, ns: non-significant, ****p < 0.0001. (D) CldU/IdU incorporation ratio from cells transfected with siControl or siRNF4 before DNA fiber labeling. Mean values (red lines). n=2, > 100 fibers per condition, ns: non-significant, ****p < 0.0001. (E) CldU/IdU incorporation ratio from cells expressing WT or the nuclease interaction dead mutant SMX* SLX4, as indicated. Mean values (red lines). n=2, > 100 fibers per condition, *p < 0.05, ****p < 0.0001. (F) CldU/IdU incorporation ratio from optoSLX4-WT cells pre-treated with Mirin (2 hr, 100 μM). Mean values (red lines). n=2, > 100 fibers per condition, ns: non-significant, ****p < 0.0001. (G) Representative images of RAD51 foci induced 60 and 120 minutes after SLX4 condensation, as indicated. Scale bar 10 μm. (H) Histogram quantification of RAD51 foci shown in (G). Data are plotted as medians. n=2, >100 cells per condition. (I) Model for the compartmentalization and activation by SLX4 of the SUMOylation/RNF4 pathway and of the structure-specific endonucleases, which leads to the extraction of TOP1-DPCs from chromatin and the degradation of nascent DNA.

The degradation of nascent DNA is expected to be initiated by SLX4-associated SSEs. We used biotin-proximity labeling to confirm the association of endogenous MUS81, EME1, XPF, and ERCC1 within optoSLX4 (Figure S6A). Consistent with this, GFP tagged MUS81, GFP tagged SLX1 and endogenous XPF colocalized with optogenetic SLX4 condensates (Figure S6B-D). To verify the contribution of the SSEs to nascent DNA degradation, we used an optoSLX4-SMX* mutant protein that is unable to interact with any of the three SSEs ^96^. Light induced compartmentalization of optoSLX4-SMX* did not induce significant degradation of CldU-labeled replication tracks (Figure 7E), despite the possible presence of endogenous SLX4 in optogenetic SLX4 compartments, which would in principle provide binding sites for SSEs. Control experiments confirmed that optoSLX4-SMX* formed light induced condensates (Figure S6E, F). If anything, the latter were more numerous and larger than wild type optoSLX4 condensates. As expected, the treatment of cells with the MRE11 inhibitor mirin also blocked the degradation of nascent DNA (Figure 7F), without affecting the formation of SLX4 condensates (Figure S6G, H, I). Furthermore, SLX4 condensation induced the progressive accumulation of RAD51 foci (Figure 7G-I). Collectively, the data indicate that SLX4 condensates promote a cascade of biochemical reactions including protein modifications with SUMO and ubiquitin and nucleolytic reactions leading to the degradation of nascent DNA and the recruitment of the recombinase RAD51 (Figure 7G, H).

## Discussion

SLX4 binds specifically to partner proteins forming protein complexes with defined stoichiometry and functional properties that can be reconstituted *in vitro* ^47, 48, 97^. Here we show that in the nucleus, the multivalent SLX4 scaffold oligomerizes extensively via SUMO-SIM and homotypic BTB interactions to locally assemble condensates in chromatin (Figure 7I). We define SLX4 foci as condensates on the basis that SLX4 molecules assemble dynamic and functional mesoscale structures that selectively concentrate proteins without a surrounding membrane and without a fixed stoichiometry. Using super resolution microscopy, we show that SLX4 condensates consist of globular clusters of approximately 100 nm-sized spherical condensates bound to chromatin. TopBP1, 53BP1 and γH2AX form similar nanometer-sized condensates that cluster together to form nuclear foci visible by conventional microscopy ^6, 66–68^. DNA topology and chromatin determine the microscopic organization of condensates assembled in response to DNA damage ^68, 98^. In addition, surface condensation on chromatin allows formation of small condensates at protein concentrations that are too low for canonical liquid-liquid phase separation ^14, 15^.

The composition of SLX4 condensates is most likely determined by the combination of SLX4 specific protein-binding interfaces and accessible SUMO conjugates that provide a platform for the recruitment of SIM-containing client proteins^70^. Using optogenetic control of SLX4 condensation with high temporal precision, we provide evidence that SLX4 condensates concentrate enzymes involved in protein modification by SUMO and ubiquitin. SLX4-mediated compartmentalization significantly increases the modification of substrate proteins. The enhancement of protein modification may result from the local concentration of enzymes and substrates within SLX4 condensates, which increases binding avidities by favoring protein rebinding after dissociation ^19^. Consistent with this, SUMOylation rates are increased up to more than 30-folds when the SUMOylation machinery is localized in the confined space of an engineered condensate ^38^.

Extensive SUMOylation of DNA replication and DNA repair proteins is required for cells to cope with DNA damage ^31, 82, 83^, and the SUMO-targeted ubiquitin ligase RNF4 promotes the necessary turnover of proteins at DNA damage sites ^83, 85, 86^. Importantly, protein SUMOylation and subsequent SUMO-targeted ubiquitylation define a replication and transcription independent pathway for the removal of topoisomerase 1, DNMT1 and PARP1 enzymes trapped on DNA by selective drugs ^87–89, 92^.

Among the protein substrates of SLX4 condensates, we identified TOP1-DPCs. SLX4 promoted the extraction of TOP1-DPCs from chromatin by activating the SUMO/RNF4 dependent protein degradation pathway. While the function of Fanconi anemia proteins in the repair of DNA interstrand crosslinks is well documented, the unexpected role of SLX4 in the processing of TOP1-DPCs demonstrated here raises the question of whether other Fanconi anemia proteins directly contribute to the repair of DNA-protein crosslinks.

Optogenetic activation of SLX4 condensates revealed that the degradation of nascent DNA also depends on the compartmentalization of the E3 SUMO ligases and the STUbL RNF4 by SLX4. This is consistent with a previous report that the collapse of stalled replication forks depends on SUMOylation and ubiquitylation of replisome components ^95^. We propose that SLX4 compartments orchestrate a cascade of enzymatic reactions in DNA repair where products become substrates for subsequent DNA repair reactions. Thus, the cellular functions of SLX4 arise from the collective behavior of proteins that make up SLX4 compartments. We found that SENP6 and RNF4 regulate the formation and the disassembly of SLX4 condensates, respectively. The SUMO-specific peptidase SENP6 controlled the level of SLX4 SUMOylation, which directly determined the efficacy of SLX4 condensation. Once formed, SLX4 condensates primed their degradation via RNF4-mediated SLX4 ubiquitylation, thereby providing a negative feedback loop.

Taken together, the data presented here suggest a mechanism for the regulated and selective modification of proteins by SUMO and ubiquitin. In this model, the local accumulation of SLX4-associated proteins within SLX4 compartments ensures the spatial confinement of protein group modifications and nucleolytic activity ^31^, while priming of condensate dissolution by SLX4 ubiquitylation regulates the confinement time of SLX4-associated enzymatic activities.

### Limitations of the study

The optogenetic system used in this study is designed to recapitulate the functions specifically arising from the assembly of SLX4 condensates, in the absence of DNA damaging agents. Consequently, the seeding of SLX4 condensates by optogenetic activation bypasses physiological mechanisms that regulate the nucleation of SLX4 condensates in response to DNA lesions and DNA replication inhibitors. For example, nucleolytic processing of replication intermediates is normally prevented by ATR, and is expected to occur only as a mechanism of last resort ^99^, yet optogenetic activation of SLX4 condensates readily enhanced the degradation of nascent DNA, in the absence of exogenous sources of DNA replication stress.

Further analysis is needed to understand the exact mode of SLX4 condensation on chromatin. We show that site-specific protein-protein interactions drive the assembly of SLX4 condensates, but we do not address directly whether SLX4 networking is associated with density transitions ^13^. We observed spherical SLX4 nanocondensates, suggesting the presence of interfacial tension ^100^, a characteristic feature of condensates formed by phase separation. Quantitative biophysical analyses of SLX4 condensates in living cells should help to determine whether SLX4 undergoes surface condensation on chromatin. It will be fascinating also to understand how SLX4 globular clusters enhance enzymatic reactions, and where exactly catalysis occurs: inside or on the surface of SLX4 nanocondensates?

## Acknowledgement

We thank Agata Smogorzewska for providing the FA-P (RA3331) cells stably transfected with expressing vector control or SLX4 expressing vector, and Ronald Hay, Stephen Jackson, Stephen West and Orian Amir for providing cDNAs. We acknowledge the national infrastructure France-BioImaging supported by the French National Research Agency (ANR-10-INBS-04). This work was supported by the Fondation ARC pour la recherche sur le cancer PGA1 RF20180206787 (A.C), the Fondation MSD AVENIR (A.C.), by the ANR BioTop project (AAPG2021) of the French Agence Nationale de la Recherche (A.C. and A.B.), by the Labex EPIGENMED, Montpellier, France (E.A.), and by the French Institut National du Cancer INCa-PLBio2016-159 and INCa-PLBio2019-152 (P-H.G.). Figure 7I was created with Biorender.com.

## Author Contributions

Conceptualization: E.A., M.P., A.B., P-H.G., J.B. and A.C.; Methodology: E.A., M.P., and J.B.; Formal analyses E.A. and M.P.; Investigation E.A., M.P., A.T., S.U. and J.B.; Writing original draft A.B. and A.C.; Writing – review and editing E.A., A.B., P-H.G., J.B. and A.C.; Visualization: E.A., M.P. and A.T.; Supervision A.B., J.B. and A.C.; Project Administration: A.C.; Funding acquisition E.A., A.B., P-H.G. and A.C.

## Inclusion and diversity

We support inclusive, diverse, and equitable conduct of research.

## STAR Methods

### Resource availability

#### Lead contact

Further information and requests for resources and reagents should be directed to the lead contact, Angelos Constantinou.

### Material availability

All unique reagents generated from this study are available upon request to the lead contact, Angelos Constantinou.

### Experimental model and subject details

#### Cell culture and transfections

Flp-In™ 293 T-REx and all stable cell lines derived from Flp-In™ 293 T-REx were grown under standard sterile cell culture conditions (37°C, 5% CO_2_, humidified incubator) in Dulbecco’s modified Eagle’s medium (DMEM) containing 10% fetal bovine serum and penicillin-streptomycin. All cells were routinely tested for mycoplasma and found negative. Parental cells were selected with 100µg/ml Zeocin and Flp-In™ 293 T-REx derived stable cell lines were maintained with 5mg/ml Blasticidin and 50mg/ml Hygromycin B.

sodium chloride (NaCl) (300 mM), sucrose and sorbitol were freshly prepared in DMEM prior to cell treatments. ML-792 (2µM), TAK-243 (2µM) or MG-132 (10µM) were added to the cell culture media for 4 hours prior to blue light exposure or Mirin (100µM) as indicated in the legend section.

For transient transfections with GFP tagged constructs (SENP6, SENP6-C1030A, RNF4, RNF4 C159A, MUS81, SLX1) cells were seeded to reach 80% on the day of transfection. For 6 well plates, 4µg of cDNA and 8uL Lipofectamine 2000 were used. Media was changed after 6 hours, and cells were analyzed for recombinant protein expression after 24 hours. For knockdown experiments, SMARTpool siRNA was purchased from Dharmacon. For each condition, a minimum of 20nM siRNA was transfected using INTERFERin transfection reagent. Knockdown efficiency was assessed 48 hours after transfection.

#### Cloning

Primers used for plasmid construction are listed in the KRT. For pcDNA5_FRT_TO_optoSLX4-wt, SIM*1,2,3 and SMX, cDNA was amplified with primers 1 and 2 using Phusion High-Fidelity DNA Polymerase. The amplified sequence was inserted into the *KpnI* site of pCDNA5_FRT_TO_TurboID-mCherry-Cry2 (Addgene 166504).

For pcDNA5_FRT_TO_mCherry-SLX4, primers 3 and 4 were used to linearize an mCherry-Cry2 construct to delete Cry2. SLX4 was amplified using primers 5 and 6. Cloning was performed using the In-Fusion HD Cloning Kit (Takara).

For the structure-function analysis, the vector pCDNA5_FRT_TO_Cry2-NLS-mCherry was prepared by amplifying Cry2 (primer 9, 10), mCherry (primer 11, 12), SLX4-NLS (primer 13, 14), and the pcDNA5 vector (primer 15, 16). Assembly was made using the In-Fusion HD Cloning Kit. This vector was linearized with PCR (primers 17, 18) and used to insert SLX4 fragments using the designated primers (primer 19-32).

Mutations in SLX4 were generated using the QuickChangeMulti Site-Directed Mutagenesis Kit. F708R with primer 33.

pcDNA5_FRT_TO_Cry2-HaloTag was generated by amplifying HaloTag with primers 36 and 37 which was inserted into the pcDNA5_FRT_TO-Cry2 vector linearized with primers 34 and 35. SLX4 was then amplified with primers 38 and 39 and inserted into the *BsiWI* site to generate pcDNA5_FRT_TO_Cry2-SLX4-HaloTag.

pcDNA3-HA-RNF4 wild-type and C159A were kindly provided by Amir Orian. RNF4 was amplified with primers 42 and 43 and inserted into pEGFP-C1 linearized with primers 40 and 41.

pBABE-eGFP-MUS81 and SLX1 were generated by cDNA amplification using primers 44-47 and inserted into the EcoRI site of the pBABE-puro-eGFP vector.

pcDNA5_FRT_TO_mCherry-F708 was generated by amplification and relegation of the original plasmid with primers 48 and 49.

### Generation of stable cell lines

Flp-In™ 293 T-REx cells are seeded to reach 80-90% confluence on the day of transfection. pcDNA5_FRT_TO expression plasmids were mixed with pOG44 encoding the Flp recombinase at a 1:7 ratio in opti-MEM. For a single transfection in a 6 well plate, 500ng of the expression plasmid was mixed with 3.5µg of pOG44 in 250uL opti-MEM. Additionally, 8uL Lipofectamine 2000 Transfection Reagent was added to 250uL opti-MEM. After an incubation period of 5 minutes at room temperature, both solutions were mixed and incubated for a further 15 minutes at room temperature. The mixture was then pipetted dropwise onto the cells. The medium was changed after 6 hours. At 48 hours post-transfection, the cells were transferred to a 100mm petri dish, and 24 hours later, the selection was performed by adding 5mg/mL Blasticidin and 50mg/mL Hygromycin B. Clones were pooled, and the cells were examined for the expression of the construct by immunoblotting and fluorescence microscopy.

### Optogenetic activation of SLX4 condensation

Cells were plated at approximately 70% confluence in DMEM. Expression of optoSLX4 was induced with 6ng/ml doxycycline for 16 hours. For light activation, the plates were transferred into a custom-made illumination box containing an array of 24 LEDs (488nm) delivering 10mW/cm2. Nucleation was induced using 3 min of light-dark cycles (4 seconds light followed by 10 seconds dark). Images were captured using a 63x objective (NA 1.46 oil). Quantification of the number of foci was performed using ImageJ (NIH).

### Live imaging

Live imaging of optoSLX4 cells was performed on a DeltaVision OMX V3/V4 microscope (GE Healthcare) equipped with a ×100/1.4 numerical aperture (NA) Plan Super Apochromat oil immersion objective (Olympus). Diode lasers at 488 and 561nm were used for cell activation and mCherry signal detection, respectively. Cells were exposed to 488nm light for 300ms, and z-stacks were acquired with an exposure time of 20ms at a frame rate of 1 frame/2 second. All images were acquired at 37°C under 5% CO2.

### Fluorescence recovery after photobleaching (FRAP)

OptoSLX4 cells were seeded into µ-Dish35 mm, high (Ibidi, 81156) and incubated for 16 hours in the presence of 6ng/ml doxycycline to induce expression of the construct. Imaging was performed with a 63x objective (NA 1.4). SLX4 condensates were photobleached and the mCherry signal intensity was measured before and for 5 minutes after bleaching. A total of 500 images were acquired during the 5-minute period. The analysis was performed using ImageJ (NIH).

### Stimulated emission depletion (STED) microscopy

Flp-In™ 293 T-REx cells expressing Cry2-SLX4-HaloTag, or Halo-SLX4 cells were stained with 200nM Janelia Fluor® HaloTag® JF646 for 15 minutes at 37°C. After 2 washes with a warm medium, cells were fixed, and processed for immunostaining with the appropriate partner antibody, and a STAR ORANGE conjugated secondary antibody (Abberior). Confocal and STED imaging was performed using a quad scanning STED microscope (Expert Line, Abberior Instruments, Germany) equipped with a PlanSuperApo 100x/1.40 oil immersion objective (Olympus, Japan). JF646 and Abberior STAR Orange were imaged at 640 and 561nm excitation. Detection was set to 650-750nm and 570-630nm, respectively. A dwell time of 10ms was used. Images were collected in line accumulation mode (5 lines), the pinhole was set to 1.0 Airy units and a pixel size of 10nm was used for all acquisitions.

### Western blot

Whole-cell extracts were obtained by lysing the cells in RIPA buffer (50mM Tris, 150mM NaCl, 1% NP-40, 1% deoxycholate, 0.1% SDS, pH 8) for 30 min on ice. After centrifugation, the supernatant was collected and the amount of protein was quantified using the Quick Start Bradford protein assay kit. Laemmli buffer was added, and proteins were boiled 5 min at 95 °C. 40μg of protein samples were resolved on precast SDS-PAGE gels (4-15% and 10%) and transferred to a nitrocellulose membrane using the Bio-Rad Trans-Blot Turbo transfer device. Membranes were blocked with 5% nonfat milk diluted in TBS-0.1% Tween 20 (TBS-T), incubated with primary antibodies overnight at 4°C, and then incubated with anti-mouse or anti-rabbit HRP secondary antibodies for 1 hour. Blots were developed with ECL according to the manufacturer’s instructions.

### Immunofluorescence staining

To visualize of optogenetically induced foci, cells expressing the designated constructs are seeded on coverslips and treated as required (inhibitors, siRNA or cDNA transfection). After light activation, the cells were fixed with PBS/4% paraformaldehyde (PFA) for 15 min at RT followed by a 5 min permeabilization and counterstaining step in PBS/ 0.2% Triton X-100/ 1mg/mL Hoechst 33342.

For immunostaining, cells grown on coverslips were fixed with PFA for 15 min at RT followed by a 10 min permeabilization step in PBS/ 0.2% Triton X-100-PBS and blocked for 30 min in PBS/3% BSA. Primary antibodies and appropriate fluorochrome-conjugated secondary antibodies were diluted in blocking solution and incubated for 1h at RT. DNA was stained with Hoechst 33342.

Coverslips were mounted on glass slides using Prolong Gold antifade reagent. Images were captured with a 63x objective (NA 1.46 oil).

For TOP1cc immunostaining, cells were seeded on coverslips. After the indicated treatment, cells were fixed with 4% PFA for 15 min on ice and then permeabilized with PBS/0.25% Triton X-100 for 15 min. Antigens were made accessible by treatment with 1% SDS for 5 min. Cells were then washed with wash buffer (PBS/0.1% BSA/0.1% Triton X-100) and blocked using 10% nonfat milk in PBS. The primary antibody (topoisomerase1-DNA cleavage complex) was incubated overnight at 1:100 in PBS containing 5% goat serum. After 5 washes with wash buffer the cells were incubated with Alexa Fluor 488-conjugated secondary antibody at 1:1000 in PBS/ 5% goat serum for 1 hour. DNA was then stained with Hoechst 33342 and coverslips were mounted with Prolong Gold antifade reagent.

### Pulldown of biotinylated proteins: TurboID

Flp-In™ 293 T-REx cell lines stably transfected with optoSLX4 recombinant protein and grown to 75% confluence were incubated with 6ng/ml of doxycycline for 16 hours. The next day, the cells were incubated with 500µM of biotin for 10 min. Cells were then washed with PBS and lysed with lysis buffer (50mM Tris-HCl pH 7.5, 150mM NaCl, 1mM EDTA, 1mM EGTA, 1% NP-40, 0.2% SDS, 0.5% sodium deoxycholate) supplemented with 1X complete protease inhibitor, 1X phosphatase inhibitor and 250U benzonase. Lysed cells were incubated on a rotating wheel for 1 hour at 4°C prior sonication on ice (40% amplitude, 3 cycles 10 sec sonication-2 sec resting). After centrifugation (7750 rcf.) for 30 min at 4°C, the cleared supernatant was transferred to a new tube and the total protein concentration was determined using the Bradford protein assay. For each condition, 2mg of proteins was incubated with 50µl of streptavidin-Agarose beads on a rotating wheel at 4°C for 3 hours. After 1min centrifugation (400 rcf.), the beads were washed sequentially with 1ml of lysis buffer, 1ml wash buffer 1 (2% SDS in H2O), 1ml wash buffer 2 (0.2% sodium deoxycholate, 1% Triton X-100, 500mM NaCl, 1mM EDTA, and 50mM HEPES pH 7.5), 1ml Wash Buffer 3 (250mM LiCl, 0.5% NP-40, 0.5% sodium deoxycholate, 1mM EDTA, 500mM NaCl and 10mM Tris pH 8) and 1ml Wash Buffer 4 (50mM Tris pH 7.5 and 50mM NaCl). Bound proteins were eluted from the agarose beads using 40µl of 2X Laemmli Sample buffer and sent for mass spectrometry analysis. For Western blot analysis of SLX4 partners enriched in optogenetic SLX4 condensates, cells were simultaneously incubated with 500µM of biotin and exposed to blue light for 10 min of light-dark cycles (4 sec light followed by 30 sec dark). Biotin proximity labeling of light induced SLX4 partners was performed using streptavidin-coated beads as described previously. Bound proteins were eluted from the agarose beads with 80µl of 2X Laemmli sample buffer and incubated at 95°C for 10 min. 5µg of the lysates were used for Western blot analysis and probed by immunoblotting to detect proteins that are associated with SLX4 clusters, in the absence of DNA damage.

### Mass spectrometry

Sample digestion was essentially performed essentially as described^101^. Briefly, proteins were loaded onto SDS-PAGE (Bio-Rad, 456-1034) and, after brief migration, a single band was excised. Proteins in the excised band were digested with trypsin (Promega). The resulting peptides were analyzed online by nano-flow HPLC-nanoelectrospray ionization using a Qexactive HFX mass spectrometer (Thermo Fisher Scientific) coupled to a nano-LC system (Thermo Fisher Scientific, U3000-RSLC). Desalting and preconcentration of samples were performed on-line on a Pepmap® precolumn (0.3 3 10mm; Fisher Scientific, 164568). A gradient consisting of 0% to 40% B in A (A: 0.1% formic acid (Fisher Scientific, A117), 6% acetonitrile (Fisher Scientific, A955), in H2O (Fisher Scientific, W6), and B: 0.1% formic acid in 80% acetonitrile) for 120 min at 300nl/min was used to elute peptides from the capillary reverse-phase column (0.075 3 250mm, Pepmap®, Fisher Scientific, 164941). Data were acquired using the Xcalibur software (version 4.0). A cycle of one full-scan mass spectrum (375–1,500 m/z) at a resolution of 60000 (at 200 m/z) followed by 12 data-dependent MS/MS spectra (at a resolution of 30000, isolation window 1.2 m/z) was repeated continuously throughout the nanoLC separation. Raw data analysis was performed using the MaxQuant software (version 1.6.10.43) with standard settings. The database used consists of Human entries from UniProt (reference proteome UniProt 2021_01) and 250 contaminants (MaxQuant contaminant database).

### Pulldown of 6xHis tagged SUMO and Ub conjugates

OptoSLX4 cells were seeded in 100mm dishes. 24 hours later, cells were transfected with 6xHis-SUMO2/3 or 6xHis-Ub. 6 hours after transfection, doxycycline was added at 6ng/ml to express optoSLX4. The next day, cells were harvested after light activation. The cell pellet was directly lysed in denaturing buffer (6M GuHCl, 0.1M NaH2PO4, 10mM Tris HCl pH 8, 5mM β-Mercaptoethanol, 5mM Imidazole). The extracts were sonicated briefly before centrifugation. The clarified extracts were incubated with TALON metal affinity resin for 1 hour at RT. Beads were washed 3 times with denaturing 8M urea buffer before elution in loading buffer supplemented with 250mM Imidazole. For experiments where knockdown was necessary, siRNA and then cDNA transfections were performed on days 2 and 3 after seeding, respectively.

### Ex Vivo/In Vitro SUMOylation Assay

Ex vivo/in vitro SUMOylation assays were performed as previously described^53^. Briefly, YFP tagged SLX4 complexes were immunoprecipitated from HeLa cell lines stably expressing YFP-SLX4 under the control of a doxycycline-inducible promoter. Cells were washed with PBS and lysed with NETN buffer (50mM Tris-HCl pH 8.0, 150mM NaCl, 1mM EDTA, 1% NP-40, 1mM DTT, 0.25mM PMSF) containing a proteasome inhibitor cocktail. After centrifugation, the supernatants were incubated with a GFP nanobody (ChromoTek GFP-Trap) for 2 hours at 4°C. The beads were washed 3 times with NETN buffer, twice with TBS and 5 times with 50mM Tris-HCl pH 8.0. The SLX4 complexes immobilized on the beads were then incubated for 60 min at 37°C in a standard reaction mixture containing 50mM Tris-HCl pH 7.6, 10mM MgCl2, 0.2mM CaCl2, 4mM ATP, 1mM DTT, 228nM of E1 (SAE1/UBA2: Boston Biochem), 2.5µM of E2 (Ubc9: Boston Biochem), and 12.5µM of SUMO2 (Boston Biochem).

### DUST assay

After exposure to the indicated treatment for the indicated time, cells were washed with ice-cold PBS and scraped in 1ml of DNAzol (Invitrogen). Nucleic acids were precipitated with 500µl isopropanol, washed with 70% ethanol, and resuspended in 300µl TE buffer. Samples were then sheared by sonication and RNA was digested with RNase A (100 μg/ml) at 37°C for 1 hour. DNA was precipitated from the samples by adding 1:10 volume of 3M sodium acetate sodium acetate and 2.5 volume of absolute ethanol. After centrifugation at 15000 rpm for 20min, DNA was resuspended in 150µl of ddH_2_O. DNA content was determined by measuring the absorbance at 260 nm (NanoDrop). 10µg of DNA per sample was digested with benzonase in the presence of 2mM MgCl2 followed by gel electrophoresis on 4-15% precast polyacrylamide gel (Bio-Rad) to detect total TOP-DPCs. For the DNA loading control and in some other experiments, samples were subjected to slot-blot for the immunodetection. Undigested DNA samples (4µg for DNA detection and 2ug for TOP1-DPCs) were resuspended in TE buffer to a final volume of 300µl per condition. A pre-soaked Whatman and a nitrocellulose membrane were placed in the slot-blot manifold setup (Heofer). The manifold wells were washed twice with TE buffer and then loaded with DNA. Each time, the vacuum was turned on only after all samples had been added to ensure even distribution. The membrane was then removed and processed as normal for immunoblotting with anti-double stranded DNA (dsDNA) and anti-TOP1.

### DNA fiber labeling

Doxycycline induced optoSLX4 cells were sequentially labeled with two halogenated thymidine analogs, 5-iodo-20-deoxyuridine (IdU at 25mM) and 5-Chloro-20-deoxyuridine (CldU at 50mM) for 30 min each. SLX4 condensation was then induced optogenetically with 3 min of light (4s) and dark (10s) cycles, every 30 min, for 2 hours. Cells were harvested and resuspended in ice-cold PBS. Two microliters of the cell suspension was applied to a microscope slide, and 7µL of spreading buffer (200mM Tris-HCl pH 7.5, 50mM EDTA, 0.5% SDS) was added on top for 3 min. DNA fibers were stretched by tilting the slide and allowing the drop to run down slowly. After fixation in a 3:1 solution of off methanol: acetic acid, the DNA was denatured with 2.5N HCl and blocked with PBS/1%BSA/0.1% Tween. DNA spreads were immunostained with mouse anti-BrdU, rat anti-CldU and mouse anti-ssDNA antibodies. Corresponding secondary antibodies conjugated to Alexa Fluor dyes were used later. Images were captured using a 40x objective (NA 1.4 oil). The acquired DNA fiber images were analyzed by using ImageJ (NIH). Statistical analysis was performed using GraphPad Prism 8. Mean values are shown as red lines. One-way ANOVA analysis was used to compare the means of samples in a group.

### Image analysis and quantification

Quantification of foci number was performed with ImageJ (NIH), using a manual pipeline based on segmentation of nuclei and evaluation of foci number by the Find Maxima function.

To measure colocalization by Pearson correlation coefficient, the EzColocalization plugin was used (Stauffer et al. 2018). A threshold algorithm was used to detect nuclei, and then a watershed segmentation was selected in the plugin to calculate the PCC value per nucleus.

### Statistical analysis and reproducibility

All statistical procedures performed are indicated in the figure legends. Each experiment was independently repeated at least three times with similar results, unless otherwise indicated.

### Molecular simulations

SLX4 molecules were represented as stickers-spacers polymers by means of a minimal model where only the BTB domain, the SIM domains, and the SUMOylation domains are represented explicitly with a one-bead-per-domain resolution. Consecutive domains were bound by harmonic bonds:

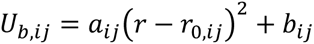

where a_ij_, r_0,ij_, and b_ij_ were parameterized to emulate the end-to-end distance distribution of the segments between the domains obtained with the Mpipi model by ^102^, which has been shown to reproduce semi-quantitatively the size of disordered proteins (see Supplemental Table 1). BTB-mediated SLX4 dimers were modeled as two monomers sharing the same BTB domain (Fig. 3C). SIM-SUMO interactions were modeled with an attractive potential previously introduced to simulate the condensation of associative biopolymers ^72^, with the following form

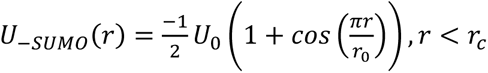

where U_0_ was set to 15 k_B_T to approximately reproduce the experimental dissociation constant for SUMO/SIM complexes of 10 µM, and r_c_ is the cut-off distance for the interaction, set to 0.8nm. Simulations were run also with U_0_ of 13 k_B_T and 17 k_B_T, corresponding respectively to K_D_ of 100 µM and 1 µM, with qualitatively similar results (Fig. S2B). All other non-bonded interactions, except for those involving deSUMOylated domains, are modeled by means of the repulsive part of the Lennard-Jones potential:

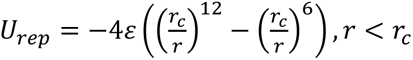

where ε is set to 1 k_B_T. The combination of these two interactions allows to impose a one-to-one valence between SIM domains and SUMO groups (Figure S3A). All non-bonded interactions with deSUMOylated groups were ignored. Non-bonded interactions between bonded particles were evaluated, to take into account intramolecular formation of SUMO/SIM complexes. All simulations were run with GROMACS 2019.6 ^103^, using a Langevin dynamics integrator with a timestep of 0.01 ps and a time constant coupling of 100 ps. In one-bead-per-domain simulations, all particles had a mass of 100 Da. Bonded and non-bonded interactions were implemented as tabulated potentials. Simulations were run in cubic boxes with periodic boundary conditions in the NVT ensemble at a temperature of 300 K. Simulations of systems with U_0_ of 15 k_B_T were run with mixtures of all possible SUMOylation patterns at constant value of SUMOylation levels. To explore the effect of the strength of SUMO/SIM interactions, we performed simulations at fixed patterns for U_0_ equal to 13, 15, and 17 k_B_T, corresponding to k_D_ of approximately 100, 10, and 1 µM, respectively, and we observed a similar dependence of SLX4 assembly on the SUMOylation degree (Figure S3B). Systems at different concentrations of SLX4 were obtained by changing the size of the simulation boxes, keeping constant the number of monomers thus maintaining the same number of SIM and SUMO particles in all simulated systems. All nucleation simulations were run with 100 monomers/50 dimers, except those with fixed patterns at varying SUMO/SIM interaction strength that were run with 500 monomers/250 dimers. Mixtures of different patterns were created inserting molecules with random patterns at a given SUMOylation level (7 combinations for level 1, 35 for level 3, 21 for level 5, and 1 for level 7) using the insert-molecules tool in GROMACS 2019.6. BTB-mediated dimers were considered as constituted by identical monomers. Simulations without intramolecular SUMO/SIM interactions where run defining for each molecule one energy group for all SIM particles and one energy group for all SUMO particles with the energygrps directive in the mdp input file, and excluding the interactions for each chain with the energygrp-excl directive in the same file. Simulations of systems with 5 SUMO sites per monomer and constant patterns extracted from single molecule simulations where run at 10 µM and 1 µM for monomers and BTB-mediated dimers, respectively. Systems at concentrations of 10 and 100 µM were simulated for 200’000’000 steps, while systems at concentrations of 1 µM were simulated for 1’000’000’000 steps due to longer equilibration times needed to observe the formation of condensed phases at low concentrations. Simulations of single molecules of SLX4 monomers and BTB-mediated dimers with all possible SUMOylation patterns were performed. The fraction of chains in the condensed phase was evaluated by considering the number of chains with at least one SUMO/SIM contact with two other chains. SUMO and SIM groups were considered in contact when their distance was below the cut-off distance r_c_. Distances between SUMO and SIM domains were evaluated by means of the gmx pairdist tool in GROMACS 2019.6 ^103^.

## Supplemental videos and Tables

Table S1: Mass spectrometric analyses of SLX4 proximal proteins, related to Figure 4A.

Video S1: Time-lapse microscopy of light-induced SLX4 condensates, image sequence n°1, related to Figure 1D.

Video S2: Time-lapse microscopy of light-induced SLX4 condensates, image sequence n°2, related to Figure 1D.

Video S3: Time-lapse microscopy of light-induced SLX4 condensates, image sequence n°3, related to Figure 1D.

## KEY RESOURCES TABLE

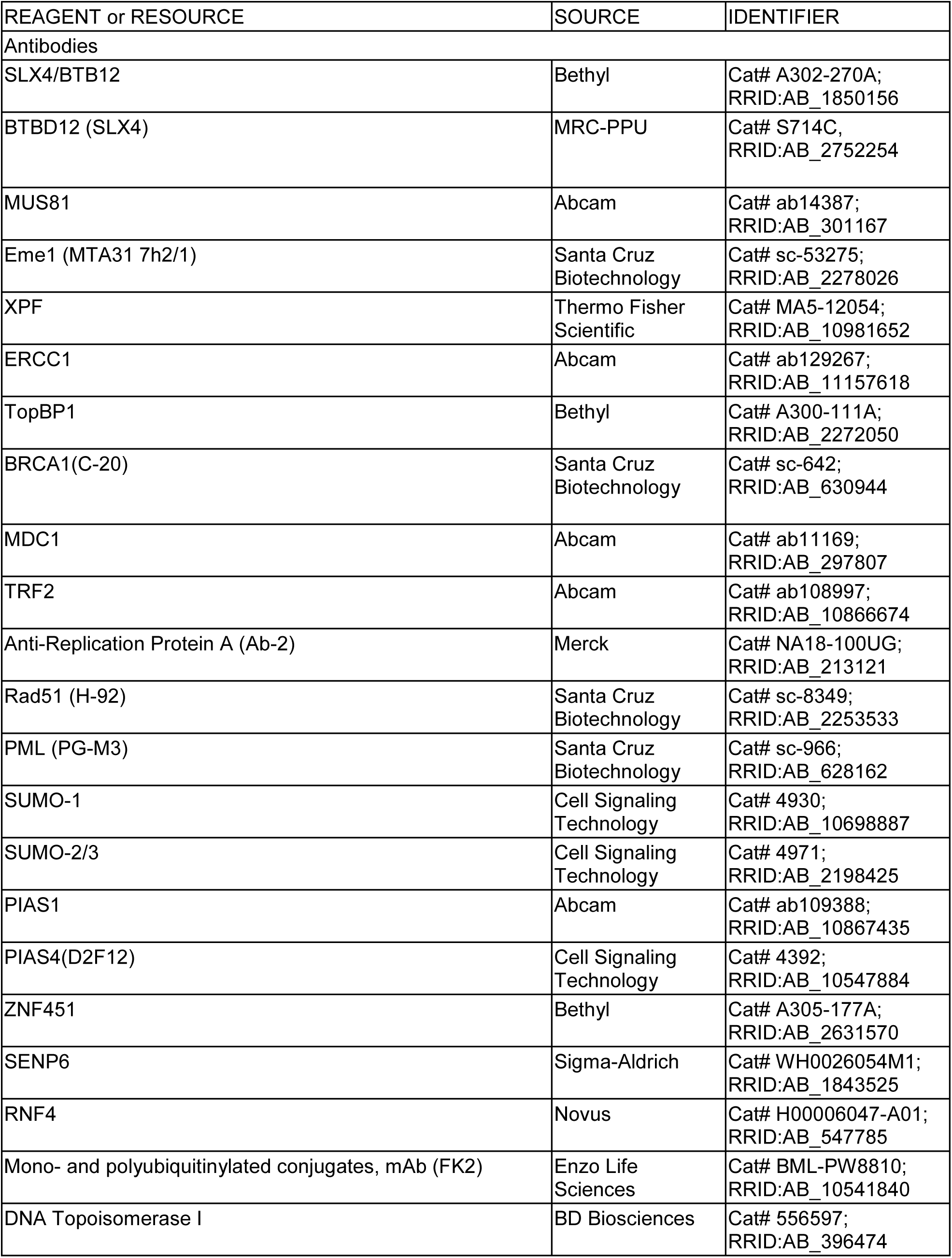
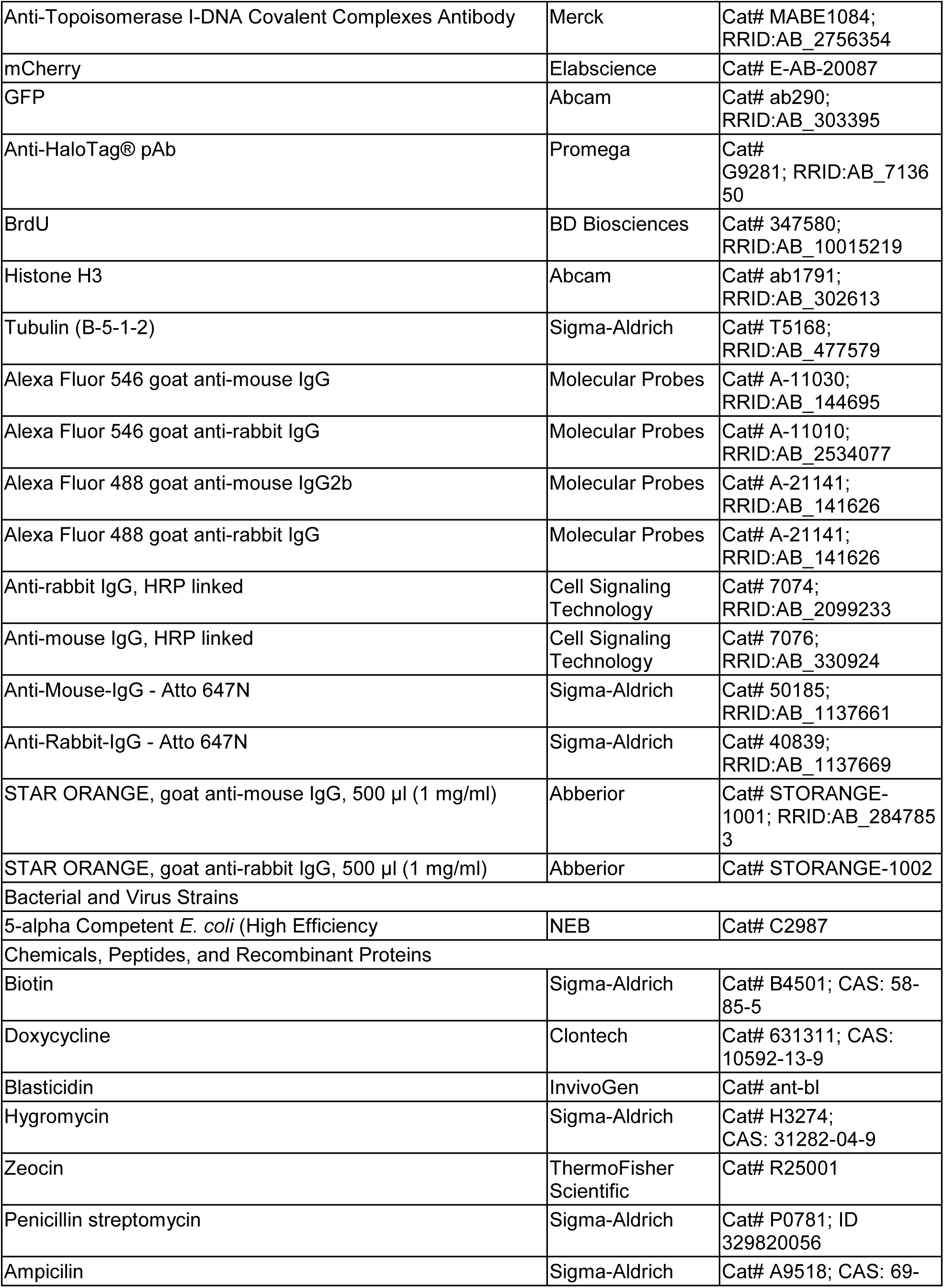
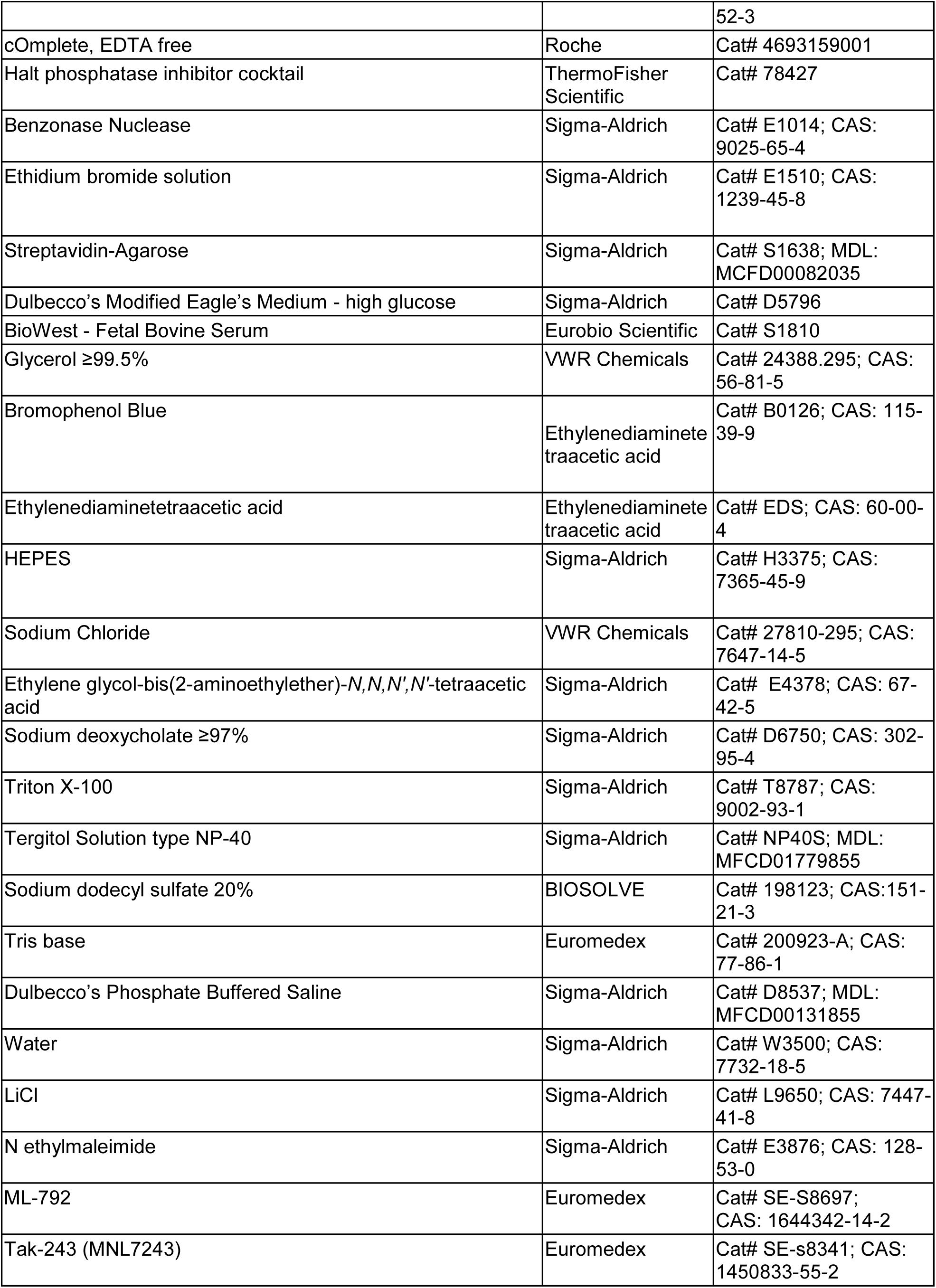
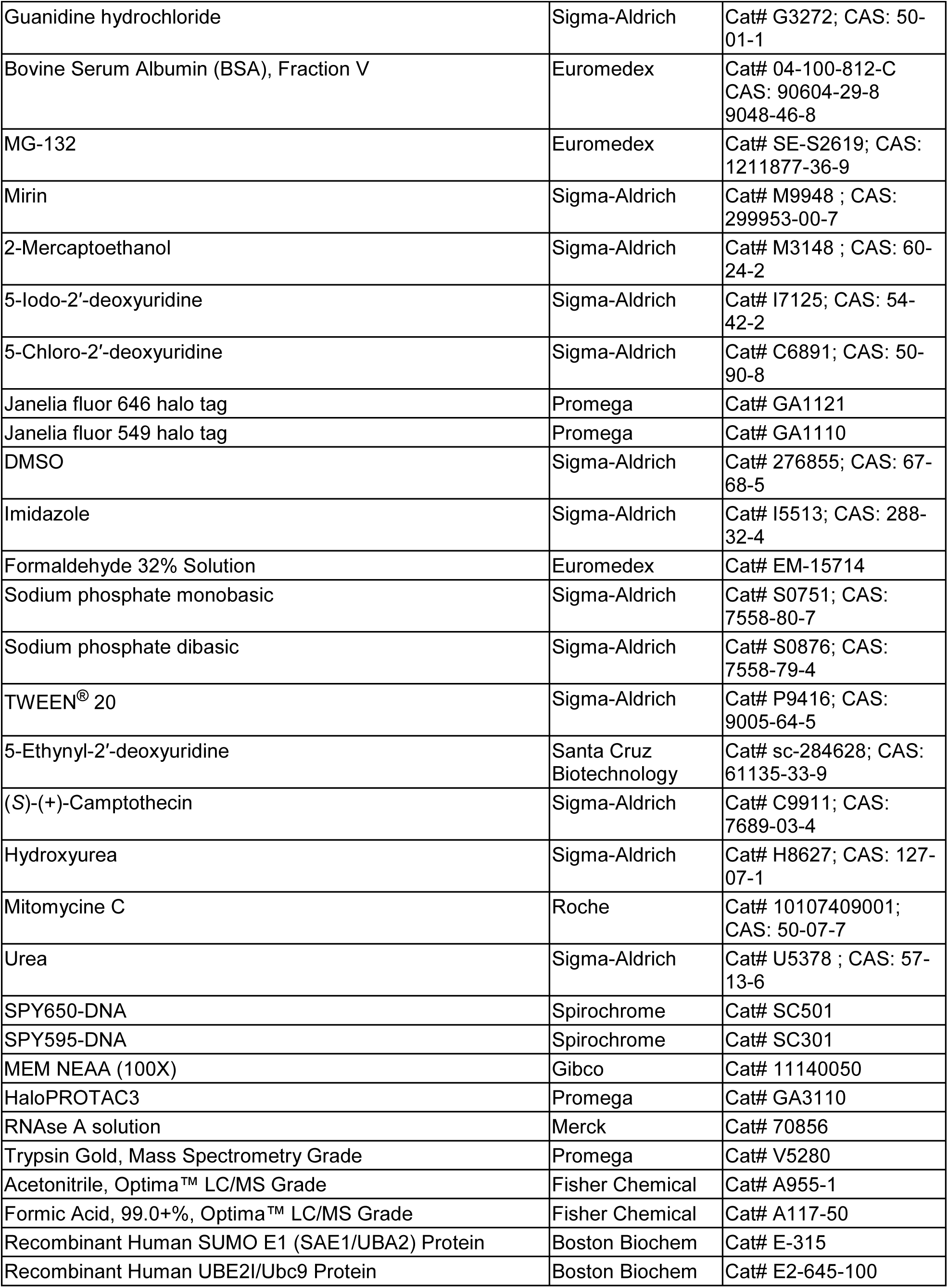
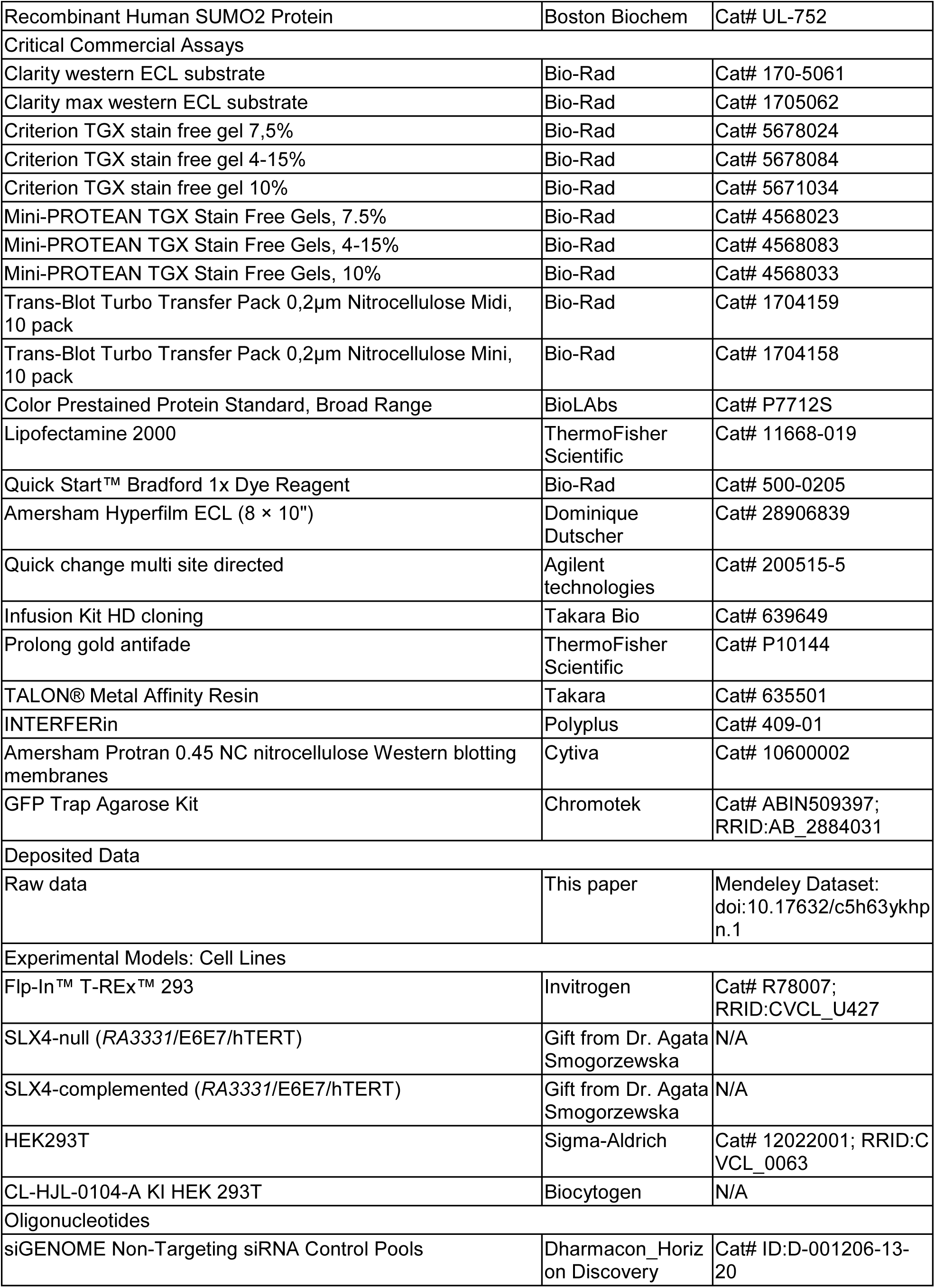
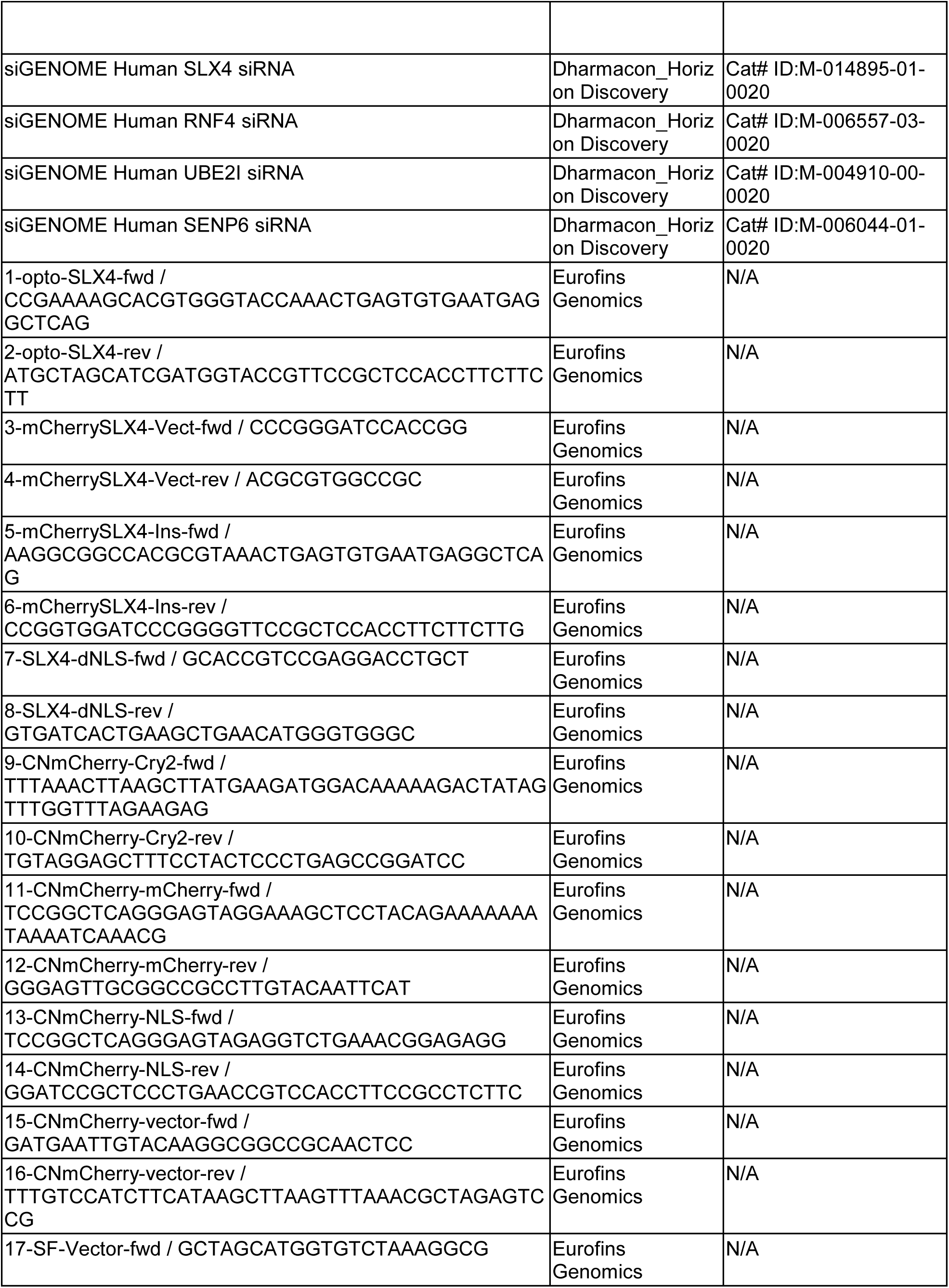
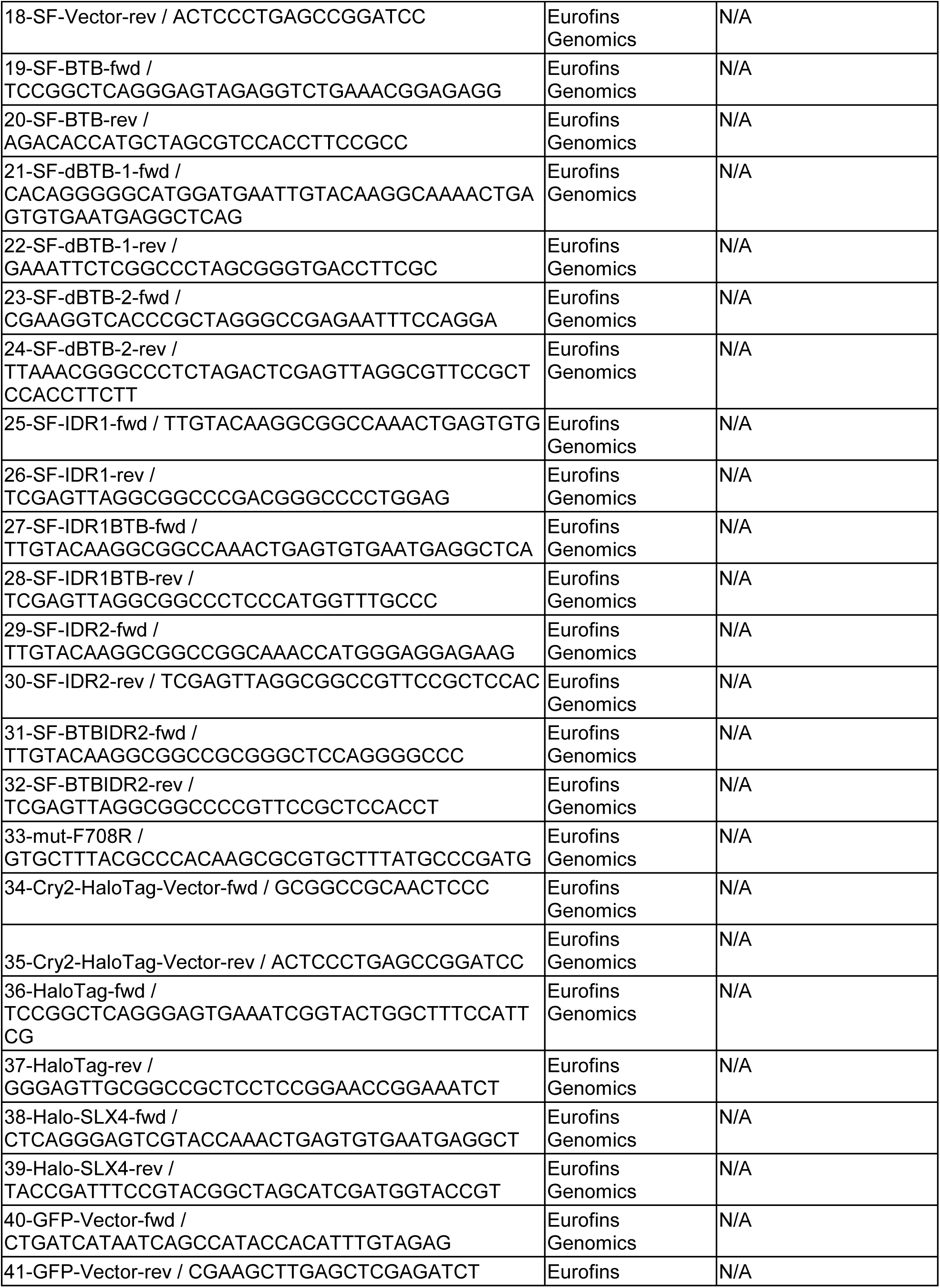
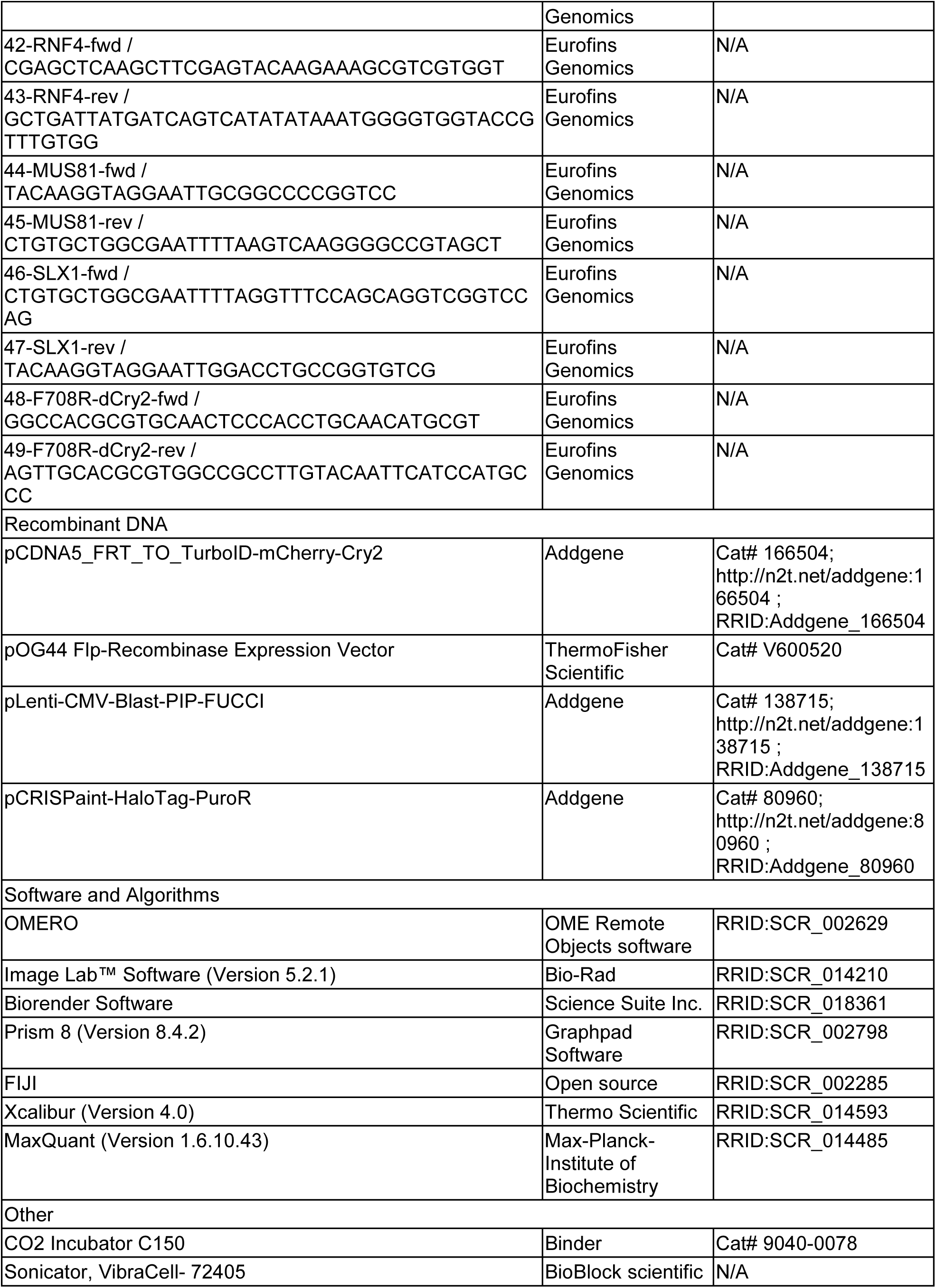
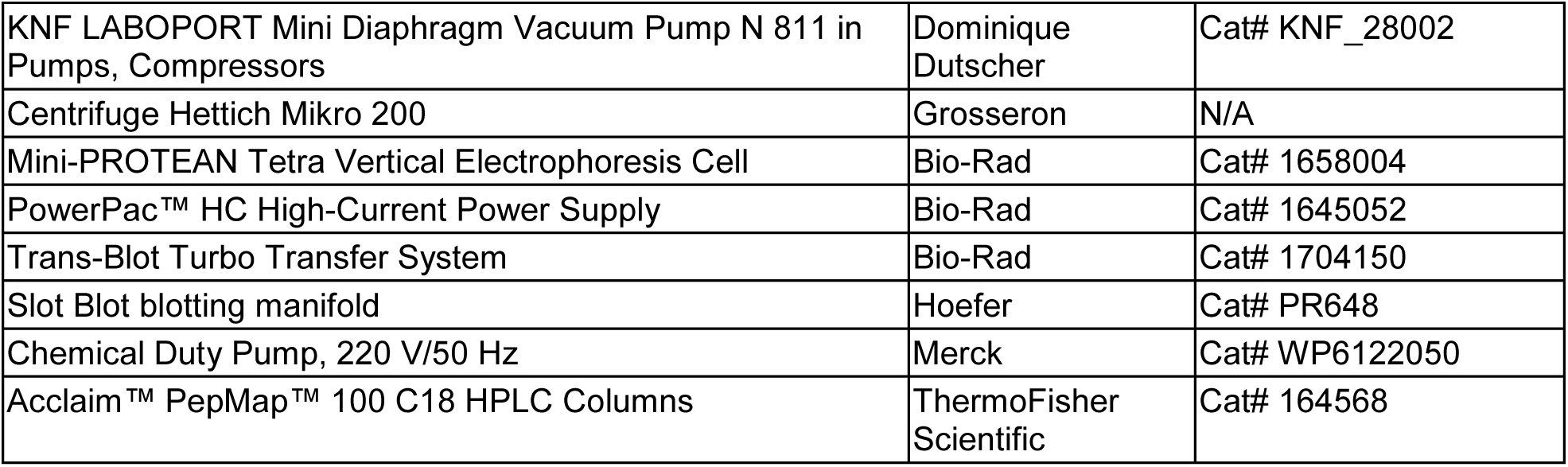

