## Supplemental Figures 1-6 and Table S1 for "Compartmentalization of the SUMO/RNF4 pathway by SLX4 drives DNA repair"

### Figure S1

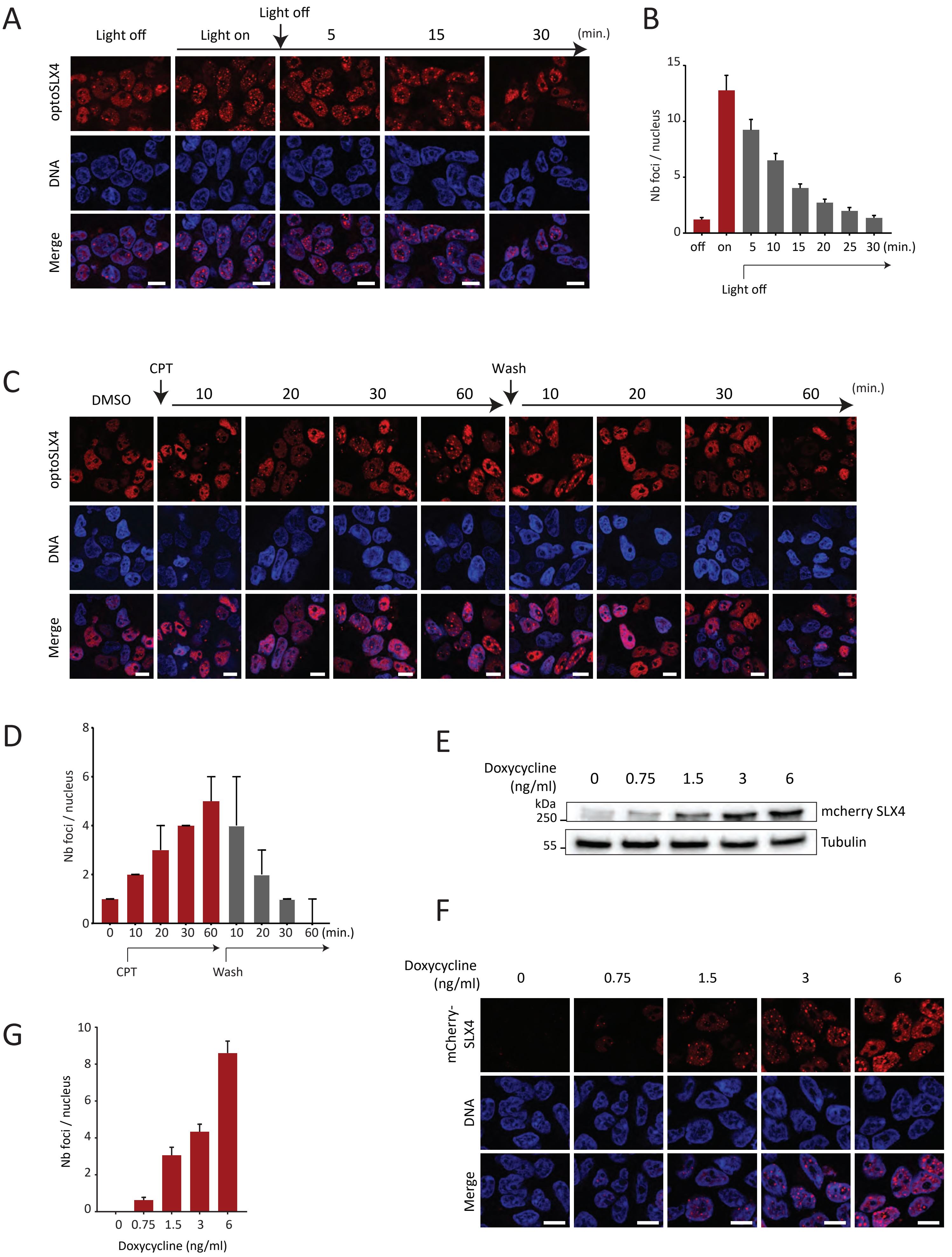

A
Light off
Light on
Light off
5
15
30
(min.)
optoSLX4
DNA
Merge
B
Nb foci / nucleus
off
on
5
10
15
20
25
30 (min.)
Light off
C
DMSO
CPT
10
20
30
60
Wash
10
20
30
60
(min.)
optoSLX4
DNA
Merge
D
Nb foci / nucleus
0
10
20
30
60
10
20
30
60 (min.)
CPT
Wash
E
Doxycycline (ng/ml)
0
0.75
1.5
3
6
kDa
250
55
mcherry SLX4
Tubulin
F
Doxycycline (ng/ml)
0
0.75
1.5
3
6
mCherry-SLX4
DNA
Merge
G
Nb foci / nucleus
0
0.75
1.5
3
6
Doxycycline (ng/ml)

**Figure S1. Characterization of SLX4 condensates. Related to figure 1**

(A) Representative images showing the dissolution of optoSLX4 condensates, 30 min after light activation. Activated cells were left in the dark for the corresponding duration. Scale bar 10 µm.

(B) Quantitative histogram analysis of SLX4 condensates in (A). Data plotted as medians with SEM; n=3, >100 cells per condition.

(C) Representative images showing the kinetics of optoSLX4 condensation under treatment with 1 µM CPT. Dissolution kinetics were monitored after washing with fresh media. Cells were collected at 10 min intervals for the indicated time. Scale bar 10 µm.

(D) Quantitative histogram of SLX4 condensates in (C). Data plotted as medians with SEM; >100 cells per condition.

(E) Immunoblotting with the indicated antibodies shows mCherry-SLX4 expression levels expressed with the designated concentration of doxycycline.

(F) Representative images of mCherry-SLX4 spontaneous foci expressed with the indicated concentration of doxycycline. Scale bar 10 µm.

(G) Quantitative histogram of spontaneous SLX4 condensates. Median (plain line), quartile (dashed line); n=2, >100 cells per condition.

Figure S2

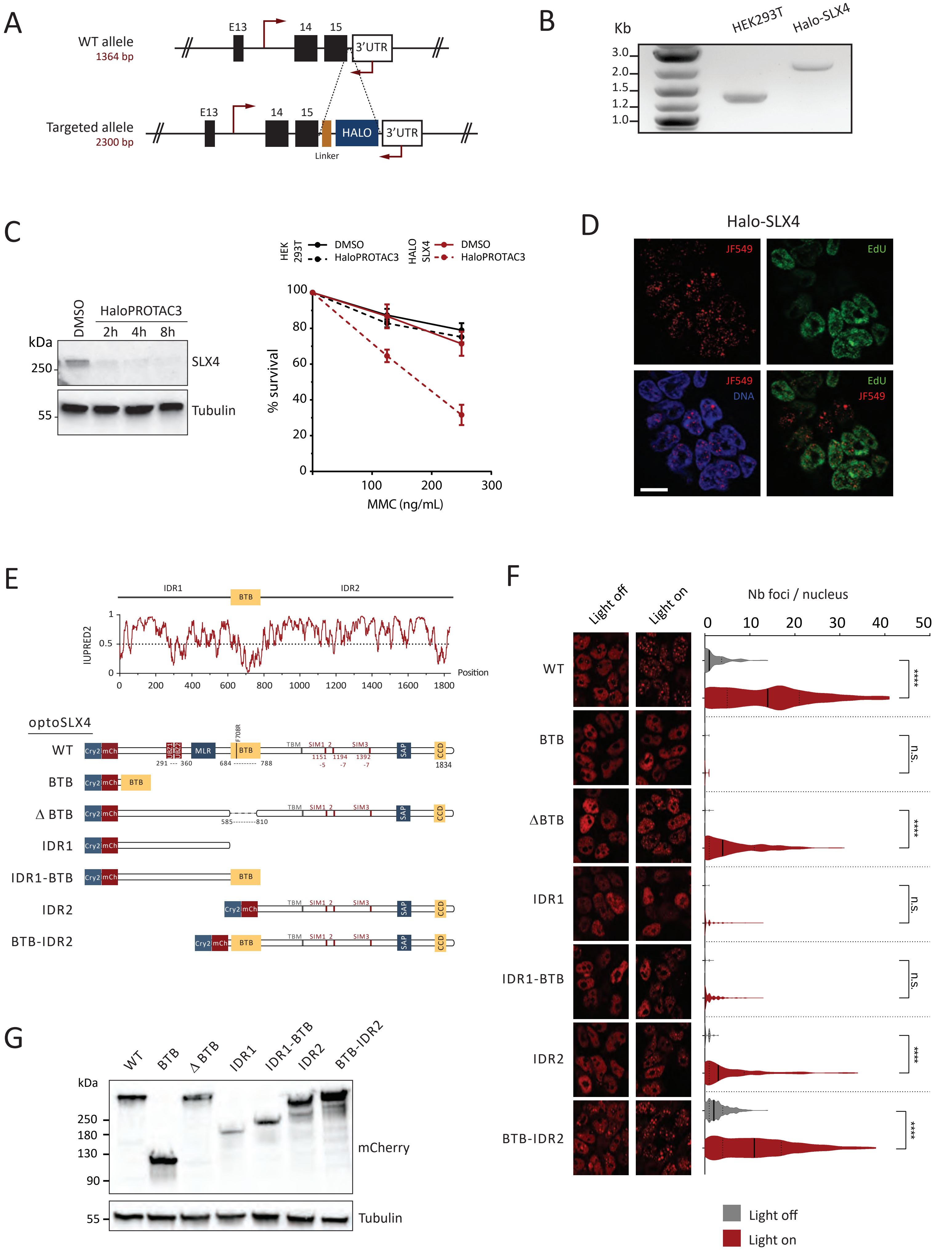
A
WT allele
1364 bp
E13
14
15
3'UTR
Targeted allele
2300 bp
HALO
Linker
B
Kb
HEK293T
Halo-SLX4
3.0
2.0
1.5
1.2
1.0
C
DMSO
HaloPROTAC3
2h
4h
8h
kDa
250
55
SLX4
Tubulin
HEK 293T
HALO SLX4
% survival
MMC (ng/mL)
D
Halo-SLX4
JF549
EdU
DNA
E
IDR1
BTB
IDR2
IUPRED2
Position
optoSLX4
WT
Cry2
mCh
MLR
TBM
SIM1
SIM3
SAP
CCD
F708R
291 --- 360
684 --------- 788
1834
ΔBTB
585---------810
IDR1-BTB
BTB-IDR2
F
Light off
Light on
Nb foci / nucleus
n.s.
****
G
mCherry
Tubulin

**Figure S2. Characterization of endogenous SLX4 foci and identification of the driving forces of SLX4 condensate formation. Related to Figures 2 and 3.**

(A) Schematic of the SLX4 knock-in strategy. The location of the genotyping primers is depicted with red arrows.

(B) Agarose gel showing the genotyping of a positive homozygous Halo-SLX4 clone compared to the parental HEK293T cells.

(C) Left, immunoblotting with the designated antibodies showing the depletion of endogenous SLX4 after treatment with HaloPROTAC3 (0.5 µM) for the indicated time. Right, quantitative measurement of colony survival of Halo-SLX4 or parental cells pre-treated with 0.5 µM HaloPROTAC3. MMC was administered with the corresponding dose for 1 hr before releasing the cells in fresh media with HaloPROTAC3 (mean ± SD, n = 2).

(D) Representative images of Halo-SLX4 spontaneous foci (red channel, JF549) and EdU incorporating cells (green channel). Scale bar 10 µm.

(E) Disorder score of human SLX4 by IUPRED2 prediction output (upper panel). Schematic diagram of the protein constructs and truncations used to investigate the domains required for SLX4 compartmentalization (lower panel).

(F) Left, representative images showing the stable cell lines expressing the different constructs in (E) before and after light induction. Right, violin plot quantification of the SLX4 foci in the corresponding cell line. Median (plain line), quartile (dashed line). n= 3, >300 cells per condition, ns: non-significant, ****p < 0.0001.

(G) Immunoblot with the indicated antibodies of the different optoSLX4 truncations.

Figure S3

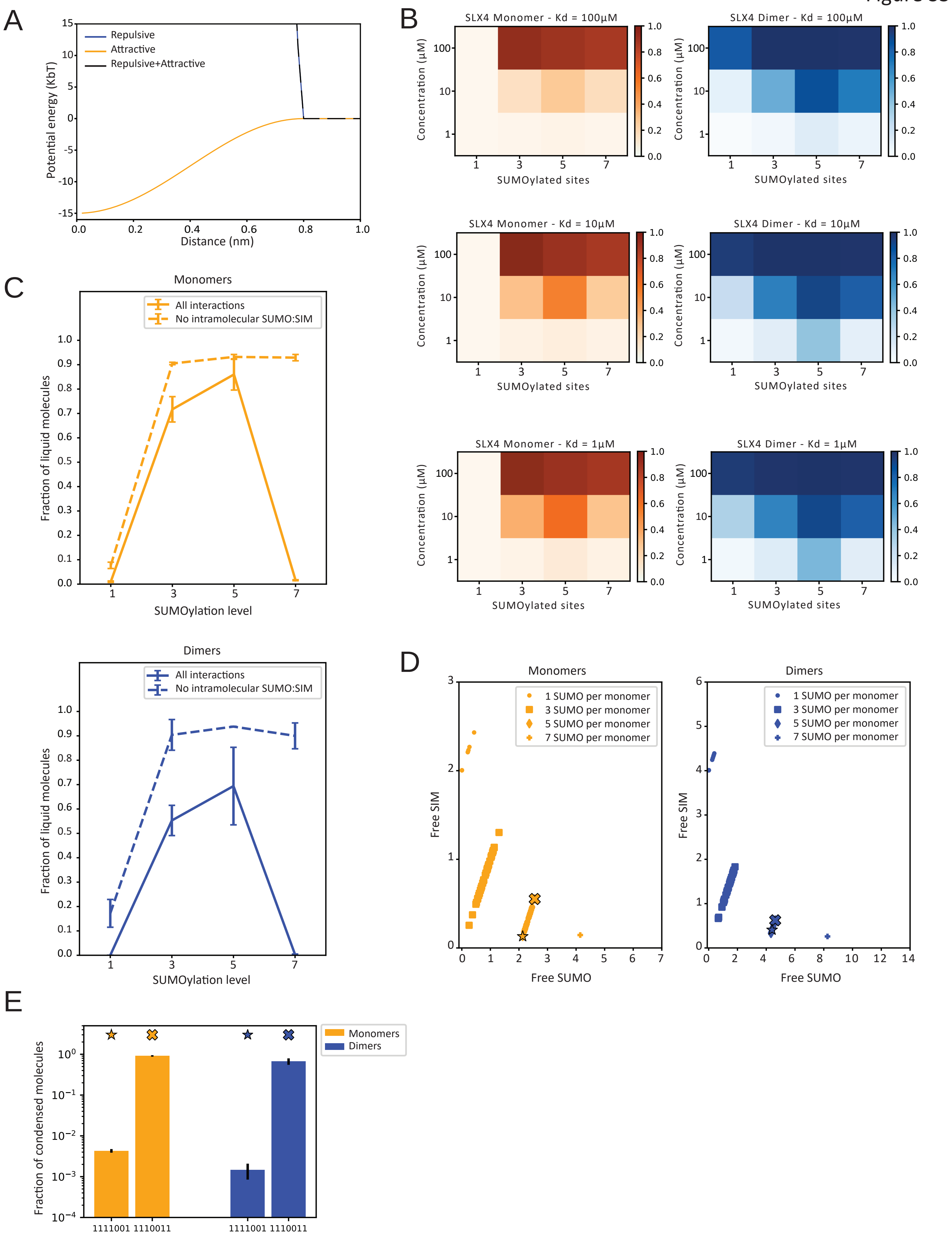
A
B
C
D
E
Repulsive
Attractive
Repulsive+Attractive
Potential energy (KbT)
Distance (nm)
SLX4 Monomer - Kd = 100µM
SLX4 Dimer - Kd = 100µM
SLX4 Monomer - Kd = 10µM
SLX4 Dimer - Kd = 10µM
SLX4 Monomer - Kd = 1µM
SLX4 Dimer - Kd = 1µM
Concentration (µM)
SUMOylated sites
Monomers
Dimers
All interactions
No intramolecular SUMO:SIM
Fraction of liquid molecules
SUMOylation level
1 SUMO per monomer
3 SUMO per monomer
5 SUMO per monomer
7 SUMO per monomer
Free SIM
Free SUMO
Fraction of condensed molecules
1111001
1110011

**Figure S3. The condensation of SLX4 depends on its dimerization and a specific patterning of SUMO interacting with SLX4 SIMs. Related to Figure 3.**

(A) Components of the non-bonded Interaction potential used in MD simulations. Homotypic interactions and interactions with BTB domains were described with a repulsive potential, while SUMO:SIM interactions were modeled with an attractive potential. The combination of the two potentials ensures that a third particle (either SIM or SUMO) is subject to an overall repulsive potential when interacting with an already formed SUMO:SIM pair, effectively limiting the interactions to a 1:1 valence.

(B) Fraction of chains in the condensed phase as a function of the total concentration and the number of SUMOylated sites per monomer in simulations of SLX4 monomers or BTB-mediated dimers using Kd values for SUMO-SIM interactions of 1 µM, 10 µM, and 100 µM, as indicated. SUMOylation patterns were (0000100), (0010101), (1110101), and (1111111) for SUMOylation levels 1,3,5, and 7 respectively. In the patterns '1' indicates a SUMOylated site, '0' a non-SUMOylated site. Different strengths of the SUMO/SIM interaction do not alter the qualitative behavior as a function of the total concentration and the SUMOylation level.

(C) Effect of removing intramolecular interactions on the propensity to form a condensed phase. Solid lines show the fraction of molecules in a condensed state as a function of the SUMOylation level for simulations of mixtures of monomers (left) and BTB-mediated dimers (right) at total concentrations of 10 µM and 1 µM, respectively, with an optimal value to form a condensed phase at a SUMOylation level of 5. When favorable intramolecular interactions are removed, the non-monotonic behavior is lost (dashed lines). Error bars are standard deviation between three independent replicas.

(D) Average number of SUMOylated sites and SIM motifs that are not involved in intramolecular interactions from single molecule simulations of all possible SUMOylation patterns for monomers (left) and BTB-mediated dimers (right) with SUMOylation levels of 1, 3, 5, and 7. For simplicity, dimers are constituted by identical monomers. Each point corresponds to a single pattern, showing that at constant SUMOylation level, patterning greatly influences the level of self-interaction. Stars and filled-crosses indicate patterns with 5 SUMO with different behaviors used to study the effect of patterning on formation of condensates (see Fig S3E). Stars are pattern (1111001) and crosses are pattern (1110011).

(E) Effect of patterning on the propensity to form condensates from simulations of monomers and BTB-mediated dimers with five SUMO per monomer in constant patterns, indicated by the labels in the x axis, at a total concentration of 10 µM and 1 µM for monomers and BTB-mediated dimers, respectively. Different patterns greatly affect the saturation concentration. Error bars are standard deviation between three independent replicas. Star and cross symbols correspond to sequences indicated with the same symbols in figure S3D.

Figure S4

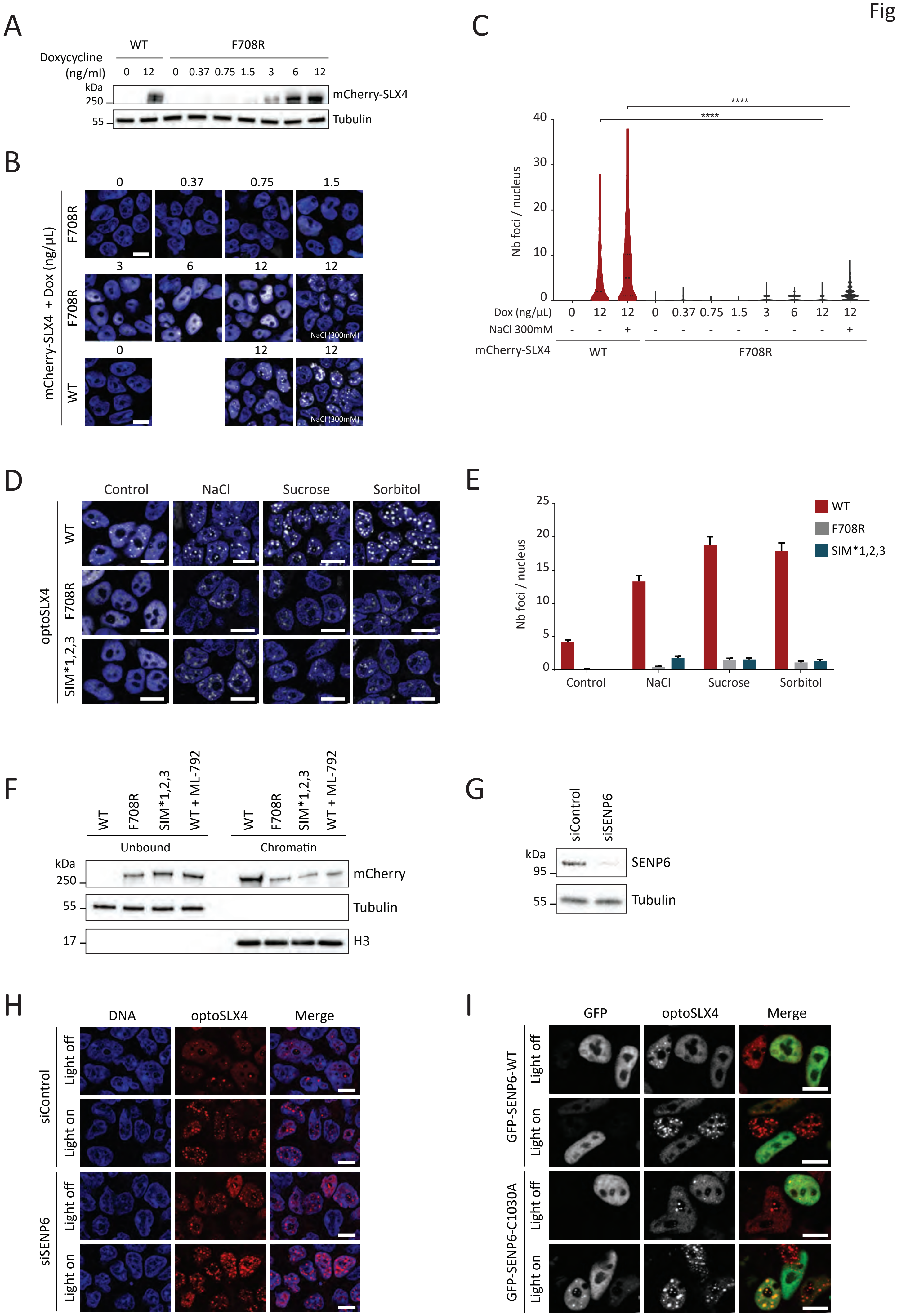

A
Doxycycline (ng/ml)
WT
F708R
0 12 0 0.37 0.75 1.5 3 6 12
kDa
250
55
mCherry-SLX4
Tubulin
B
mCherry-SLX4 + Dox (ng/µL)
F708R
WT
0 0.37 0.75 1.5
3 6 12 12
0 12 12
NaCl (300mM)
C
Nb foci / nucleus
****
40
30
20
10
0
Dox (ng/µL) 0 12 12 0 0.37 0.75 1.5 3 6 12 12
NaCl 300mM - - + - - - - - - - +
mCherry-SLX4 WT F708R
D
Control NaCl Sucrose Sorbitol
optoSLX4
WT
F708R
SIM*1,2,3
E
Nb foci / nucleus
25
20
15
10
5
0
Control NaCl Sucrose Sorbitol
WT
F708R
SIM*1,2,3
F
WT F708R SIM*1,2,3 WT + ML-792
Unbound
Chromatin
kDa
250
55
17
mCherry
Tubulin
H3
G
siControl siSENP6
kDa
95
55
SENP6
Tubulin
H
DNA optoSLX4 Merge
siControl
siSENP6
Light off
Light on
I
GFP optoSLX4 Merge
GFP-SENP6-WT
GFP-SENP6-C1030A
Light off
Light on

**Figure S4. The BTB and SUMO:SIM interactions drive SLX4 condensate formation on chromatin. Related to Figure 4.**

(A) Immunoblotting with the indicated antibodies shows the levels of mCherry-SLX4-WT and F708R expressed with the designated concentration of doxycycline.

(B) Representative images of mCherry-SLX4-WT and F708R spontaneous foci expressed with the indicated concentration of doxycycline. Cells were subjected to osmotic stress with 300mM NaCl when specified. Scale bar 10 µm.

(C) Violin plot quantification of spontaneous SLX4 foci in (B). Median (plain line), quartile (dashed line); >400 cells per condition, ****p < 0.0001.

(D) Representative images of optoSLX4-WT, F708R and SIM*1,2,3 expressing cells after being subjected to osmotic stress for 10 min (300mM NaCl, sucrose, or sorbitol). Scale bar 10 µm.

(E) Quantitative histogram analysis of SLX4 condensates in (D). Data plotted as medians with SEM; n=2, >100 cells per condition.

(F) Immunoblotting with the indicated antibodies of fractionated cell extracts prepared from cells expressing optoSLX4-WT, F708R and SIM*1,2,3 treated with doxycycline. When specified, optoSLX4 cells were incubated for 2 hr with 2 µM of ML-792.

(G) Immunoblotting for the indicated proteins from cells treated with siControl or siSENP6.

(H) Representative images of optoSLX4 cells transfected with the indicated siRNAs prior to light activation. Scale bar 10 µm.

(I) Representative images of optoSLX4 cells transfected with GFP-SENP6-WT or GFP-SENP6-C1030A, as indicated, before and after light activation. Scale bar 10 µm.

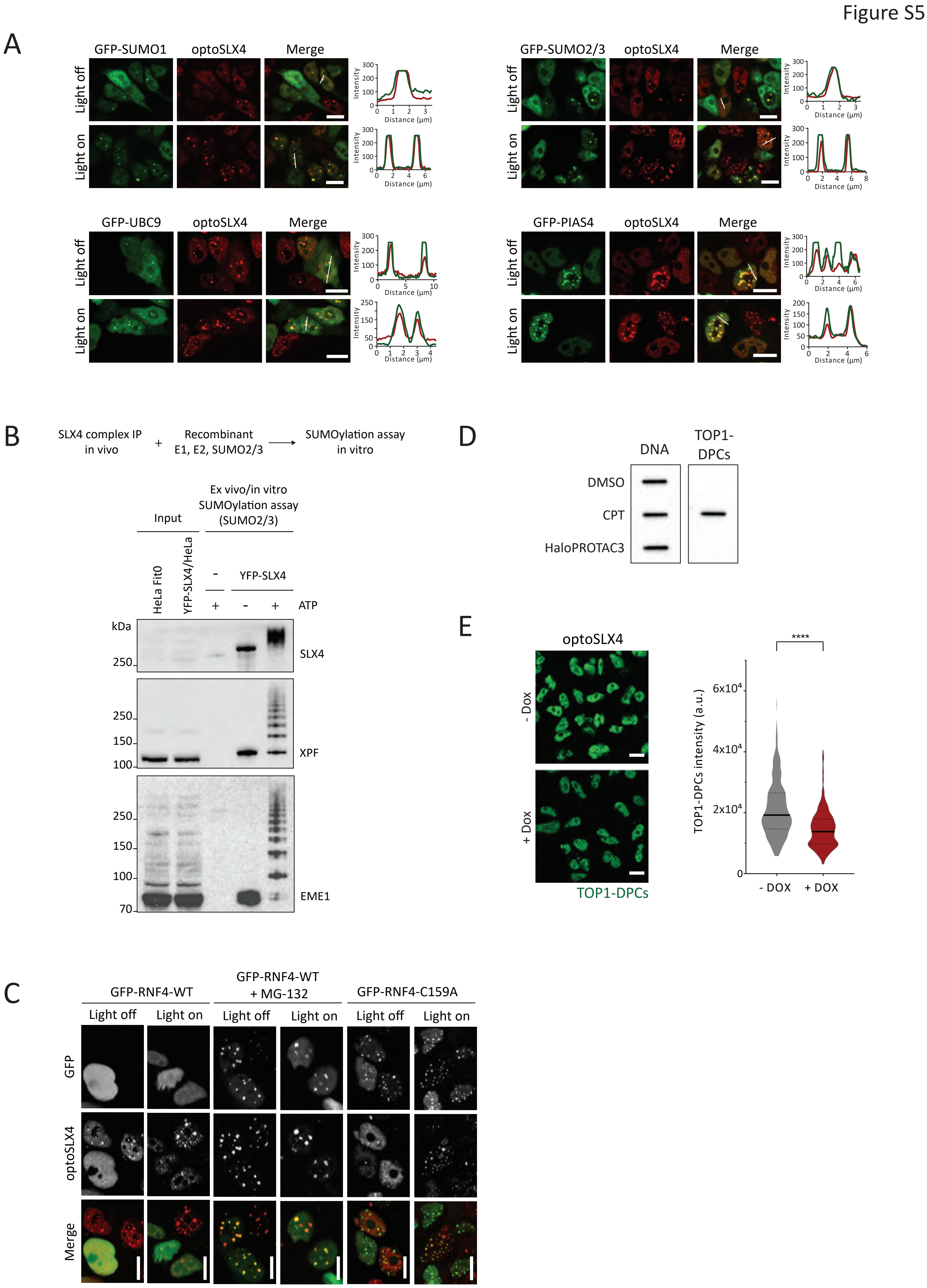
Figure S5
A
GFP-SUMO1
optoSLX4
Merge
Light off
Light on
Intensity
Distance (µm)
GFP-SUMO2/3
GFP-UBC9
GFP-PIAS4
B
SLX4 complex IP in vivo
+
Recombinant E1, E2, SUMO2/3
SUMOylation assay in vitro
Input
Ex vivo/in vitro SUMOylation assay (SUMO2/3)
HeLa Fit0
YFP-SLX4/HeLa
-
YFP-SLX4
+
-
+
ATP
kDa
250
150
100
70
SLX4
XPF
EME1
C
GFP-RNF4-WT
GFP-RNF4-WT + MG-132
GFP-RNF4-C159A
Light off
Light on
GFP
optoSLX4
Merge
D
DNA
TOP1-DPCs
DMSO
CPT
HaloPROTAC3
E
optoSLX4
- Dox
+ Dox
TOP1-DPCs
****
TOP1-DPCs intensity (a.u.)
6x10^4
4x10^4
2x10^4
0
- DOX
+ DOX

**Figure S5. SLX4 condensates recruit the SUMOylation machinery and are controlled by RNF4. Related to Figures 4, 5 and 6.**

(A) Representative images of optoSLX4 cells (red channel) transfected with GFP tagged SUMO1, SUMO2/3, UBC9 and PIAS4 (green channel), before and after light activation. Line scans of the red and green channels show co-localization. Scale bar 10 µm.

(B) Upper panel, Schematic diagram for the ex vivo/in vitro SUMOylation assay. Lower panel, immunoblotting with the indicated antibodies following the SUMOylation assay.

(C) Representative images of optoSLX4 cells transfected with GFP-RNF4-WT or GFP-RNF4-C159A, as indicated, before and after light activation. When specified, cells were treated with MG132 (10 µM) for 4 hr. Scale bar 10 µm.

(D) Slot blot of TOP1-DPCs and DNA purified from HEK293T cells under chaotropic conditions following treatment with 1 µM CPT or 0.5µM HaloPROTAC3 for 2 hr. DNA was used as a loading control.

(E) Left, representative images showing the level of TOP1-DPCs in doxycycline induced versus un-induced optoSLX4 cells treated with 1 µM CPT for 15 min. Scale bar 10µm. Right, violin plot quantification of TOP1-DPCs intensity. Median (plain line), quartile (dashed line); n= 2, >200 cells per condition, ****$p < 0.0001$.

Figure S6

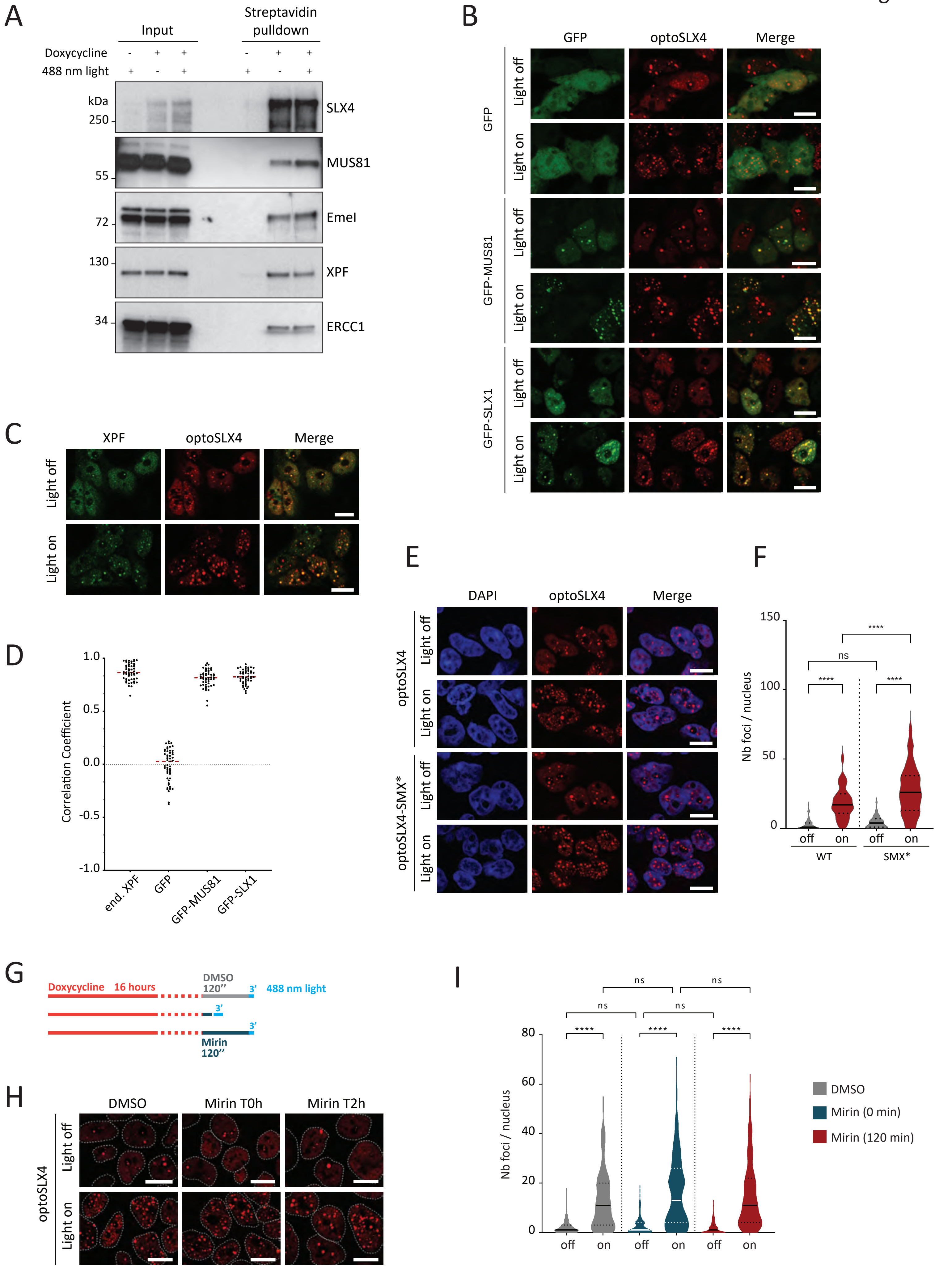

A
Input
Streptavidin pulldown
Doxycycline
488 nm light
kDa
250
55
72
130
34
SLX4
MUS81
EmeI
XPF
ERCC1
B
GFP
optoSLX4
Merge
Light off
Light on
GFP-MUS81
GFP-SLX1
C
XPF
D
Correlation Coefficient
1.0
0.5
0.0
-0.5
-1.0
end. XPF
E
DAPI
optoSLX4-SMX*
F
Nb foci / nucleus
150
100
50
0
ns
****
off
on
WT
SMX*
G
Doxycycline 16 hours
DMSO
120''
3'
488 nm light
Mirin
H
DMSO
Mirin T0h
Mirin T2h
I
80
60
40
20
DMSO
Mirin (0 min)
Mirin (120 min)

**Figure S6. SLX4 condensates recruit the structure specific endonucleases. Related to Figure 7.**

(A) Immunoblotting for the indicated proteins isolated with streptavidin beads from optoSLX4 cells treated with doxycycline and exposed to 488 nm light, when specified (+). Biotin was added to the media in all conditions.

(B) Representative images of optoSLX4 cells transfected with GFP-control and GFP-tagged MUS81 and SLX1, before and after light activation. Scale bar 10 µm.

(C) Representative images of optoSLX4 condensates and endogenous XPF revealed by immunofluorescence staining. Scale bar 10 µm.

(D) Quantification, using Pearson's correlation coefficient, of the co-localization between the indicated proteins and optoSLX4 condensates. Median shown in red, >100 cells per condition.

(E) Representative images of optoSLX4-WT and SMX* before and after exposure to 488 nm light. Scale bar 10 µm.

(F) Violin plot quantification of (E). Median (plain line), quartile (dashed line); >200 cells per condition, ns: non-significant, ****p < 0.0001.

(G) Schematic diagram of the Mirin treatment to assess its effect on SLX4 condensation.

(H) Representative images of optoSLX4-WT cells treated with Mirin as indicate in (G) before and after exposure to 488 nm light. Scale bar 10 µm.

(I) Violin plot quantification of (H). Median (plain line), quartile (dashed line); >200 cells per condition, ns: non-significant, ****p < 0.0001.

**Table S1. Parameters for the bonded interactions between the explicit domains in the minimal model of SLX4. Related to Figure 3 and Figure S3.**

The first column reports the residue numbers of the SLX4 sequence of the first and last amino acids of the protein segment modeled with a harmonic bond. Note that residues between 684 and 788 correspond to the sequence of the BTB domain.

| **SLX4 residues** | **$x_{0,ij}$ [nm]** | **$a_{ij}$ [kJ/mol/nm$^2$]** | **$b_{ij}$ [kJ/mol]** |
|---|---|---|---|
| 291-684 | 11.5475 | 0.0754653 | 9.82136 |
| 788-902 | 7.76986 | 0.158887 | 9.12396 |
| 902-1151 | 11.2579 | 0.0604855 | 10.5146 |
| 1151-1169 | 2.43396 | 2.7647 | 5.0439 |
| 1169-1179 | 1.55051 | 11.3416 | 2.78733 |
| 1179-1194 | 2.10861 | 4.21843 | 4.45981 |
| 1194-1392 | 9.35967 | 0.0779045 | 10.0876 |
| 1392-1575 | 9.26161 | 0.0886052 | 10.0053 |
| 1575-1588 | 1.95533 | 5.71225 | 3.86846 |
| 1588-1657 | 5.3133 | 0.316272 | 8.21493 |
